# Zebrafish Foxl2l suppresses stemness of germline progenitors and directs feminization

**DOI:** 10.1101/2024.04.03.587938

**Authors:** Chen-wei Hsu, Hao Ho, Ching-Hsin Yang, Yan-wei Wang, Ker-Chau Li, Bon-chu Chung

## Abstract

Zebrafish is an important organism for genetic studies, but its germ cell types and the mechanism of sex differentiation remain elusive. Here, we conducted a single-cell transcriptomic profiling and charted a developmental trajectory going from germline stem cells, through early, committed, and late progenitors, to pre-meiotic and meiotic cells. A transcription factor, Foxl2l, is expressed in the progenitors committed to the ovary fate. CRISPR-Cas9-mediated mutation of *foxl2l* produced 100% male fish with normal fertility. Another single-cell profiling of *foxl2l^-/-^* germ cells reveals the arrest of early progenitors. Concomitantly the expression of *nanos2* (stem cell marker) and *id1* (transcription repressor in stem cells) was elevated together with an increase of *nanos2^+^* germ cell in *foxl2l* mutants, indicating the reversion to the stem cell state. Thus, we have identified developmental stages of germ cells in juvenile zebrafish and demonstrated that Foxl2l drives zebrafish germ cell progenitors toward feminization and prevents them from reverting back to the stem cell state.

## Introduction

Sex differentiation is an important yet complicated subject. The mammalian sex is determined by the XX/XY system, in which the *SRY* gene on the Y-chromosome plays a pivotal role (Berta et al., 1990b). Females lack *SRY*. They rely on such genes as RSPO1, *FOXL2* and *WNT4* to promote female differentiation and to antagonize male differentiation (She & Yang, 2017). In contrast to higher vertebrates, fishes have more complicated schemes of sex differentiation. Wild zebrafish strain Nadia uses the female-dominant ZZ/ZW system, but domesticated laboratory strains have no sex chromosome (Wilson et al., 2014). Furthermore, male and female zebrafish are morphologically indistinguishable at the larval stage presenting an obstacle in the study of their sex differentiation. Zebrafish oocytes start to develop at around 21 days post fertilization (dpf) (Tong et al., 2010), while male differentiation appears later at 28 dpf (Luzio et al., 2021). The sex-differentiation mechanism in zebrafish is unclear.

Germ cell abundance and the existence of oocytes play essential roles in the differentiation of zebrafish gonad. Zebrafish containing reduced numbers of germ cells (*nr0b1* mutants) or no germ cell (*dnd* morphans) develop into males (Chen et al., 2016; Slanchev et al., 2005). Mutations of genes (*figla*, *sycp3* or *bmp15*) involved in oocyte development also result in female-to-male sex reversal (Dranow et al., 2016; Pan et al., 2022; Qin et al., 2018). These results indicate that intrinsic signals from germ cells are required for the female fate. However, the developmental stages of germ cells in zebrafish remain unclear.

One transcription factor that promotes female differentiation is Foxl2l (forkhead box L2-like). Medaka (*Oryzias latipes*) *foxl2l*, named *foxl3* (Liu et al., 2022), encodes a germline intrinsic factor that suppresses spermatogenesis and initiates oogenesis in XX gonad (Kikuchi et al., 2020). XX medaka fish with *foxl3* mutation become hermaphroditic possessing functional sperms (Nishimura et al., 2015). Zebrafish *foxl2l* is a marker of the pre-meiotic progenitor germ cells. Its disruption leads to 100% male fish (Liu et al., 2022), signifying the role of Foxl2l in female development. However, the mechanism of Foxl2l action remains poorly understood.

In this study, we conducted single-cell RNA sequencing (scRNAseq) analysis to dissect the developmental trajectory of germ cells in zebrafish during the critical sex determination stage. In addition to the known germline stem cells and meiotic germ cells, we have further identified three subpopulations of progenitor cells, early progenitor, committed progenitor and late progenitor. We conducted another scRNAseq of *foxl2l* mutant zebrafish and proved that Foxl2l controls the development of committed progenitor, which is indispensable for female differentiation. We further show that Foxl2l suppresses the reversion of progenitors to the stem cell state.

## Results

### Developmental trajectory of zebrafish germ cells

Germ cells play important roles in zebrafish sex differentiation, but their development is poorly studied. We set out to investigate their development during the initial stage of sex differentiation at 26 days post fertilization (dpf) (Luzio et al., 2021; Uchida et al., 2002). Fluorescent germ cells were isolated from *piwil1:EGFP* transgenic fish by cell sorting (Supplementary Figure S1) for single-cell RNA sequencing (scRNAseq) using the 10X Genomics platform (Figure 1A). After data preprocessing, we obtained single-cell transcriptomes for clustering analysis. The germ cells were partitioned into 14 clusters and visualized on UMAP (Figure 1B). Trajectory analysis identified a linear structure for germ cell development, and pseudotime analysis ordered these clusters as W1 to W14 (Figure 1C). Marker genes from each cluster were found and their expression profiles were displayed in a heatmap (Supplementary Figure S2). Their expressions appear to stay high in the neighboring clusters, exhibiting a successive pattern of. We also showed the expression profiles of the top 37 out of a total of 1491 marker genes on UMAP separately (Supplementary Figure S3).

**Fig. 1.**
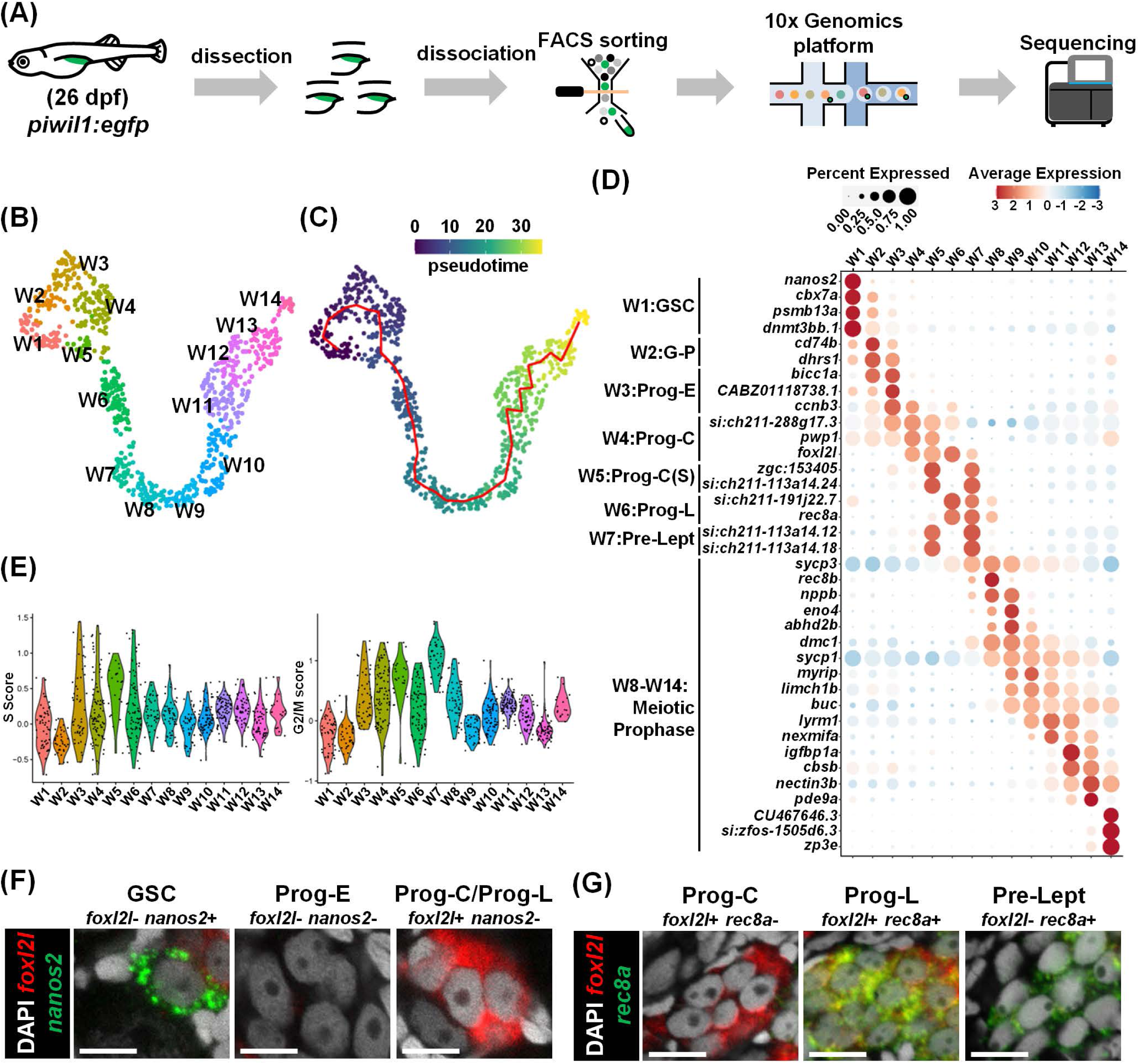
Single cell transcriptome landscape of zebrafish germ cell development. **(A)** Flow chart for sample collection and RNA sequencing. **(B)** WT germ cells in 14 clusters visualized by *Uniform Manifold Approximation and Projection* (UMAP). **(C)** The pseudotime shown in color bar and the trajectory shown by a red line. **(D)** Expression of top marker genes for each cluster in a dot plot. **(E)** The S and G2/M cell cycle scores in each cluster shown by violin plots. **(F,G)** Identification of germ cell types by RNA fluorescent *in situ* hybridization (FISH) of marker genes, *nanos2* and *foxl2l* in (F), *rec8a* and *foxl2l* in (G), in 26-dpf gonads. Scale bar: 10 µm.

Using known markers we annotated zebrafish germ cells to include three broad categories, germline stem cells (GSC), progenitors (Prog), and meiotic germ cells. Cells in cluster W1 were identified as GSC because they express *nanos2* (Figure 1D) (Beer & Draper, 2013; Z. Cao et al., 2019). Clusters W3-W6 were identified as progenitors because they have high cell cycle scores (Figure 1E), and progenitors are known to divide synchronously forming germline cysts (Bertho et al., 2021). Cluster W2 lies between W1 and W3 with reduced *nanos2* expression, indicating they transit from GSC to progenitor (G-P). Cells in clusters W8 to W14 are enriched with meiosis expressing meiotic markers (*sycp1*, *sycp3*, *dmc1*, *buc* and *zp3e*) (Gautier et al., 2013)(Pan et al., 2022)(Marlow & Mullins, 2008; Wassarman & Litscher, 2021; Yoshida et al., 1998). Cluster W7 lies between progenitors and meiotic cells; it has a high S-phase score typical of Pre-leptotene (Pro-Lept) at the premeiotic S-phase (Figure 1E).

A portion of progenitors (W4-W6) expressed the progenitor gene *foxl2l* (Figure 1D), suggesting them as progenitor subtypes (Liu et al., 2022). We defined cluster W3 as early progenitors (Prog-E) for the absence of *foxl2l*, cluster W4 as committed progenitors (Prog-C) for the expression of *foxl2l*, cluster W5 as Prog-C(S) for the expression of both *foxl2l* and typical S-phase genes, *zgc:153405* and *si:ch211-113a14.24* (orthologs of linker histone *H1* genes), and cluster W6 as the late progenitor (Prog-L) for the expression of *foxl2l* and genes for meiotic entry such as *rec8a* (ortholog of *REC8*) and *si:ch211-191j22.7* (ortholog of *MEIOSIN*, the meiosis initiator) (Figure 1D).

In addition to our own data, we also extracted scRNAseq data of 40-dpf female germ cells from a public database (Liu et al., 2022). Following the same data analysis procedure, the developmental trajectory and marker gene expression profiles were found consistent with ours (Supplementary Figure S4), suggesting the validity of our data and the finer developmental transitions of germ cells in juvenile gonad as charted above.

### Histological identification of progenitor germ cells

Zebrafish progenitors are round in shape with one to three nucleoli at the center of the nucleus, and morphologically indistinguishable from GSC (Tong et al., 2010). To characterize these three progenitor subpopulations, we examined the marker gene expression using double fluorescence *in situ* hybridization (FISH) and detected GSC as positive for *nanos2*. Prog-E was negative for both *nanos2* and *foxl2l,* whereas both Prog-C and Prog-L were *foxl2l^+^nanos2^-^*(Figure 1F). Double FISH also detected Prog-C as *foxl2l^+^ rec8a^-^*, Prog-L as *foxl2l^+^ rec8a^+^*, and pre-leptotene cells as *foxl2l^-^rec8a^+^* (Figure 1G).

### Expression of *foxl2l* becomes female biased during sexual differentiation

We examined temporal expression of *foxl2l* by *in situ* hybridization during the sexual differentiation period. We used the Nadia strain for its defined ZZ/ZW (male/female) sex chromosomes, facilitating the distinction of their sexes at the juvenile stage (Wilson et al., 2014). The *foxl2l* expression was similar between ZZ and ZW gonads at 11 dpf before apparent sex differentiation (Figure 2A). Upon female differentiation at 22 dpf, *foxl2l* expression was decreased in the male ZZ gonad. The female-biased expression became more obvious at 30 dpf. In adult ovary at 5 months post fertilization (mpf), *foxl2l* was highly expressed in cystic germ cells but was barely observed in testis.

**Fig. 2.**
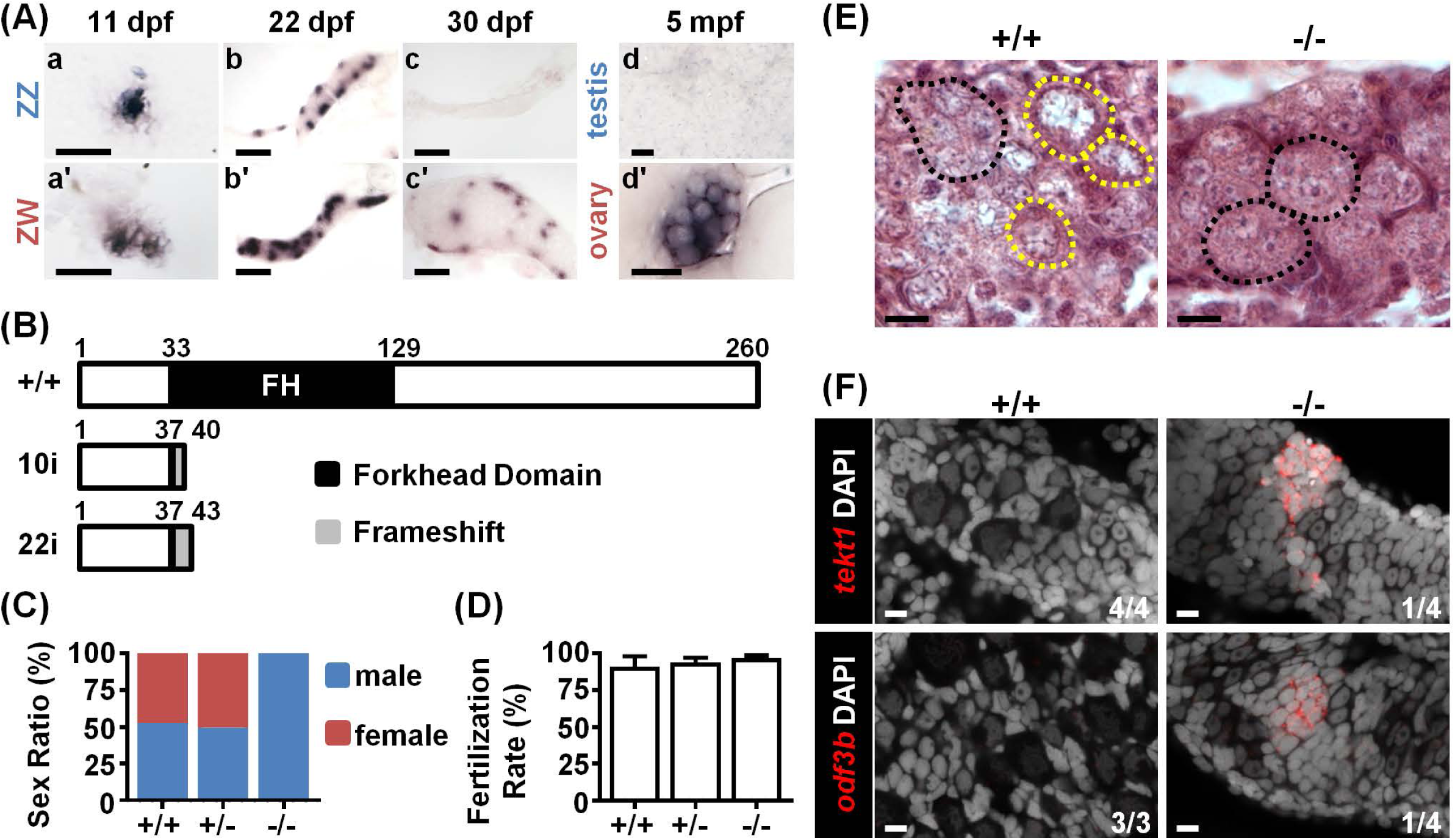
Foxl2l is required for female differentiation. **(A)** *In situ* hybridization detects *foxl2l* transcripts in bipotential gonads at 11 dpf (a, a’). Expression of *foxl2l* becomes female biased when female differentiation initiates at 22 dpf (b, b’), and becomes female-specific at the initiation of male differentiation at 30 dpf (c, c’). In adults, *foxl2l* is expressed in cystic cell in ovary but not in testis (d, d’). mpf: months post fertilization. All samples were collected from Nadia strain. Scale bars are 50 µm in (a, a’), 100 µm in (b, b’, c, c’), 20 µm in (d, d’). **(B)** Domain structure of WT and the predicted mutant of Foxl2l with 10 base pairs insertion (10i) or 22 base pairs insertion (22i) generated by CRISPR/Cas9 system. **(C)** Sex ratios of WT (+/+), *foxl2l^+/22i^* heterozygous (+/-) and *foxl2l^22i/22i^*homozygous (-/-) mutants at 4 months of age reveal that *foxl2l* mutant fish are all males. **(D)** All *foxl2l^10i/10i^* mutant males (-/-) are fertile. **(E)** The *foxl2^22i/22i^*homozygous (-/-) mutants lack meiotic oocytes at 20 dpf shown by histological staining. Black dashed circle: germ cell cysts with one to three nucleoli in each cell. Yellow dashed circle: meiotic oocyte. Scale bars represent 20 µm. **(F)** The *foxl2l^22i/22i^* mutants (-/-) express male markers (*tekt1* or *odf3b*) detected by RNA FISH at 28 dpf. Scale bars are 10 µm.

### All *foxl2l* mutants fail to enter female meiosis and become fertile males

To investigate the function of *foxl2l*, we generated *foxl2l* mutants using CRISPR-Cas9. Two resultant mutant lines, *foxl2l^10i/10i^*and *foxl2l^22i/22i^*, contained DNA insertions (Supplementary Figure S5A, S5B) causing a premature stop codon disrupting the FH domain of Foxl2l (Figure 2B). All *foxl2l* mutants became males (Figure 2C), and all *foxl2l* mutant males were fertile (Figure 2D).

We further examined histology of mutant gonads. At 20 dpf, diplotene oocytes with large perinucleolar nuclei were observed in WT gonads, but only non-meiotic cystic germ cells were found in *foxl2l* mutant (Figure 2E). At 28 dpf, small and round germ cells that express spermatogenic markers, *tekt1* and *odf3b* (Nishimura et al., 2015), were found in mutant but not WT gonads (Figure 2F). Therefore, in *foxl2l* mutants, germ cells fail to enter female meiosis, and develop into fertile males. This indicates that *foxl2l* is required for female but not for male development.

### *foxl2l* mutant germ cells are arrested at committed progenitor

We examined transcriptomic profiles of germ cells from 26-dpf *foxl2l* homozygous mutants using scRNAseq, and assigned developmental stages to them using WT marker genes at each developmental stage (Supplementary Figure S6A). The expression of the 37 top WT marker genes in mutant cells was displayed on UMAP (Supplementary Figure S6B). We found that these marker genes were expressed at similar stages as they represented for WT cells.

Pie charts showed that most of the mutant cells were annotated as GSC, G-P, Prog-E or Prog-C, with only a few as Prog-C(S), the stage before Prog-L differentiation (Figure 3A). None of the mutant cells were at Prog-L or meiotic stages (Figure 3A). This indicates that mutant germ cells cannot progress beyond Prog-C. To verify this statement, we used fluorescent staining and detected Prog-L marker *rec8a* in cystic germ cells and meiotic marker Sycp3 in meiotic cells of the control (Figure 3B). Both *rec8a* and Sycp3 staining were missing in *foxl2l* mutant gonads. These data show that mutant germ cells do not enter the state of Prog-L.

**Fig. 3.**
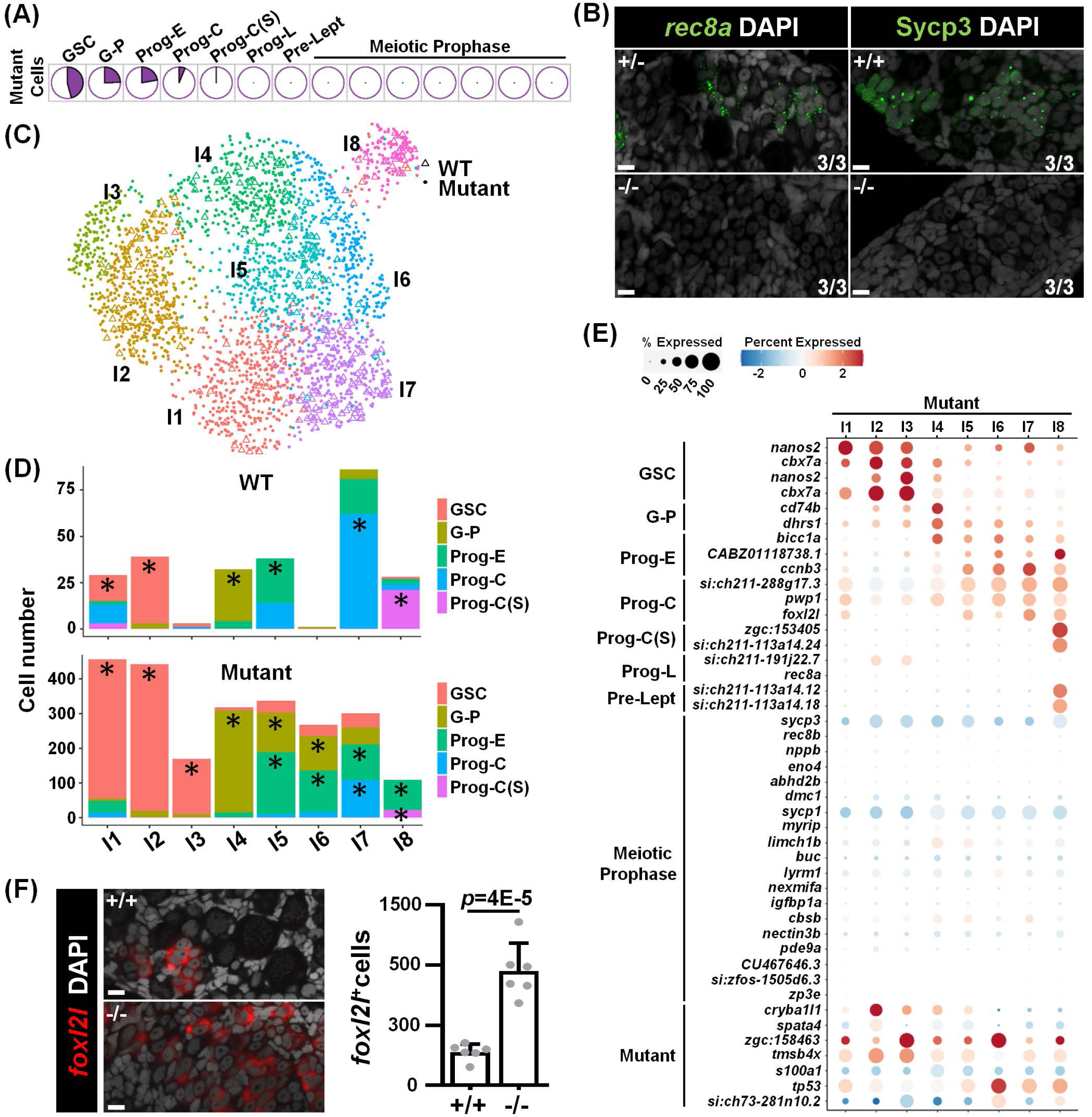
Germ cells in *foxl2l* mutant are arrested at the stage of committed progenitor with aberrant gene expression. **(A)** The proportion of mutant cells classified into different developmental stages in pie charts. **(B)** Absence of Prog-L marker, *rec8a*, and meiotic marker, Sycp3, in *foxl2l* mutant gonads. (Left) RNA FISH and (Right) immunofluorescence staining of WT (+/+), *foxl2l^+/10i^* (+/-), and *foxl2l^10i/10i^* (-/-) gonads at 28 dpf (Left) and 21 dpf (Right), respectively. N=3 in each genotype. Scale bars represent 10 µm. **(C)** Grouping of integrated cells in 8 clusters visualized by UMAP. Hollow triangle: WT cell. Solid dot: mutant cells. **(D)** The proportions of developmental stages for WT and mutant cells in each integrated group shown as bar graphs. Dominant stages with statistical significance were marked by asterisks (*). **(E)** Expression of marker genes (Y-axis) in the mutant cell in the integrated groups (X-axis) in a dot plot. **(F)** The *foxl2l^10i/10i^* (-/-) gonads contain increased number of *foxl2l^+^* cells at 26 dpf. Left: *In situ* hybridization of the gonads with *foxl2l*. Right: quantification of the staining data. One dot in the graph represents one gonad.

The component of WT cells from GSC to Prog-C stages were zoomed in for closer examination (Supplementary Figure S7A). The data of these cells were used for co-analysis with mutant germ cells.

### Altered developmental program in *foxl2l* mutant

The mutant germ cells were heterogeneous encompassing diverse cell types. In order to understand the different subtypes of mutant cells and their relationship with the WT cells, we co-clustered all mutant cells with WT cells from GSC to Prog-C stages in a scheme shown in Supplementary Figure S7B. This process resulted in eight integrated groups (I1-I8) (Figure 3C). WT and mutant cells with similar transcriptomic profiles were put in the same integrated group.

We examined the distribution of cells at different stages across different integrated groups (Figure 3D). Dominant stages were obtained after calibration against their prevalence rates in the entire cell population. For WT cells, we found WT GSC dominant in groups I1 and I2, G-P dominant in group I4, Prog-E dominant in group I5, Prog-C dominant in group I7, and Prog-C(S) dominant in group I8 (Figure 3D).

For mutant cells, while groups I5-I8 had two dominant stages, the dominant stages in each groups are similar to those in WT (Figure 3D). This indicates that WT and mutant cells with high cell-to-cell similarity belong to similar developmental stage. Groups I1 and I2 were dominated by GSC, but the expression patterns of top GSC marker genes were slightly different with higher expression of *nanos2* in I1 but of *cbx7a*, *psmb13a* and *dnmt3bb.1* in I2 (Supplementary Figure S7C). This suggests that GSCs can be further divided into two finer stages, GSCI and GSCII.

Groups I3 and I6 are particularly unique. They contained few WT cells, and were classified as GSC for I3 and as Prog-E for I6 mutant cells (Figure 3D). I3 mutant cells additionally expressed *zgc:158463* and I6 mutant cells expressed *tp53* and *zgc:158463* (Figure 3E). This indicates that I3 and I6 mutant cells expressed different genes from WT GSCs and progenitors, potentially leading to a unique cell fate.

Furthermore, the Prog-C marker *foxl2l* was expressed in mutant cells of I1, I5, I7, and I8, which were mostly classified as GSC or Prog-E in addition to Prog-C (Figure 3E). The number of *foxl2l^+^*cells detected by FISH was also increased in mutant gonads compared to WT (Figure 3F). The aberrant *foxl2l* expression in some earlier GSC and Prog-E plus the accumulation of *foxl2l*-expressing cells in the mutant indicate that progenitor development is impaired in *foxl2l* mutants.

### Differential gene expression between WT and *foxl2l* mutant germ cells

We further evaluated the effect of *foxl2l* mutation on gene expression throughout the development. Stratified by integrated groups, we compared the WT and mutant cells at the same dominant stages within each integrated group, which is defined as a stage-matched set. The stage-matched cells, color coded GSCI, GSCII, G-P, Prog-E, Prog-C and Prog-C(S) are shown in Figure 4A. Figure 4B shows genes that were differentially expressed between WT and mutants. We found that Prog-C(S) contained the largest number of genes with high expression fold-changes, consistent with the failure of mutant cells at this stage to differentiate further to Prog-L (Figure 4B). The expression of genes such as *stk31* was decreased and that of *ptmab* was increased upon transition to Prog-C in the WT cells; yet these genes were steadily expressed in the mutant progenitors (Supplementary Figure S8). This pinpoints the defect in *foxl2l* mutant germ cells during the transition from Prog-E to Prog-C.

**Fig. 4.**
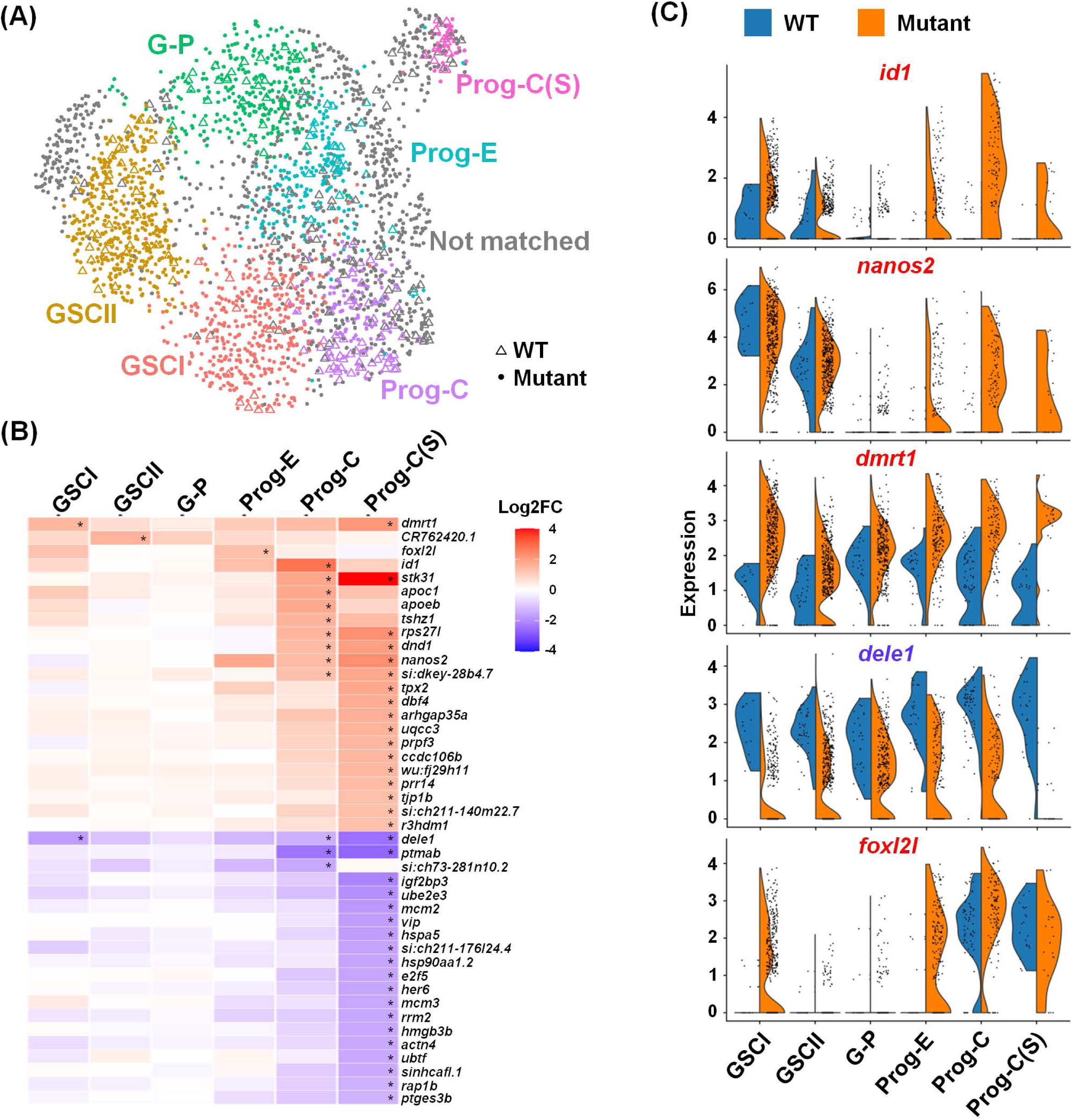
Identification of differentially expressed genes (DEGs) in WT and *foxl2l* mutants. **(A)** Visualization of the integrated cells by UMAP. Six matched stages of WT and mutant cells are shown in different colors. Cells that are not matched are labeled in gray. Hollow triangle: WT cells. Solid dot: mutant cells. **(B)** Heatmap showing fold change of gene expression between the mutant versus WT at each stage. Developmental stages are shown in the X-axis, and gene names are shown in the Y-axis. Asterisks indicate genes with significant gene expression difference (fold change (FC) of mutant/WT ≥ 2.5, *q-*value ≤ 0.05, and fraction ≥ 0.5). **(C)** Split-violin plots showing the distribution of cells expressing top DEGs at different development stages (X-axis) between WT and mutant.

The top DEGs, *dmrt1* and *id1*, and the GSC marker *nanos2* were further analyzed (Figure 4C). The *id1* and *nanos2* transcripts were low in the WT progenitors but became abundant in mutant Prog-C and Prog-E (Figure 4C). The expression of *dmrt1* and *dele1* in WT was steady throughout the developmental stages (Figure 4C), but *dmrt1* was abnormally high and *dele1* was abnormally low in both mutant GSCI and Prog-C/Prog-C(S). The *foxl2l* transcript was also unexpectedly detected in mutant GSCI and Prog-E (Figure 4C). These results indicate multiple defects of gene expression in the mutants.

### Co-expression of *foxl2l* with *dmrt1*, *nanos2* or *id1* in *foxl2l* mutant

The expression patterns of *foxl2l* along with *nanos2*, *id1* and *dmrt1* at the early stages were further examined. The *nanos2* and *foxl2l* genes were distinctly expressed among WT cells, but co-expressed in some mutant cells (Figure 5A). Double FISH also detected cells expressing both *nanos2* and *foxl2l*, and the proportion of *nanos2^+^foxl2l^+^*cells among *foxl2l^+^* cells was increased in the mutant (Figure 5B). Moreover, the number of *nanos2^+^* cell increased in mutants at 21 dpf during the early stage of female differentiation (Figure 5C). To further investigate whether GSC is affected in mutants, we counted the number of *nanos2^+^foxl2l^-^* cells, and found that the number increased dramatically in mutants at 26 dpf (Figure 5D). The co-expression of GSC marker, *nanos2*, in the *foxl2l*-expressing Prog-C and the increase of GSC cells in mutant together suggest that some Prog-C may revert to GSC in the mutant.

**Fig. 5.**
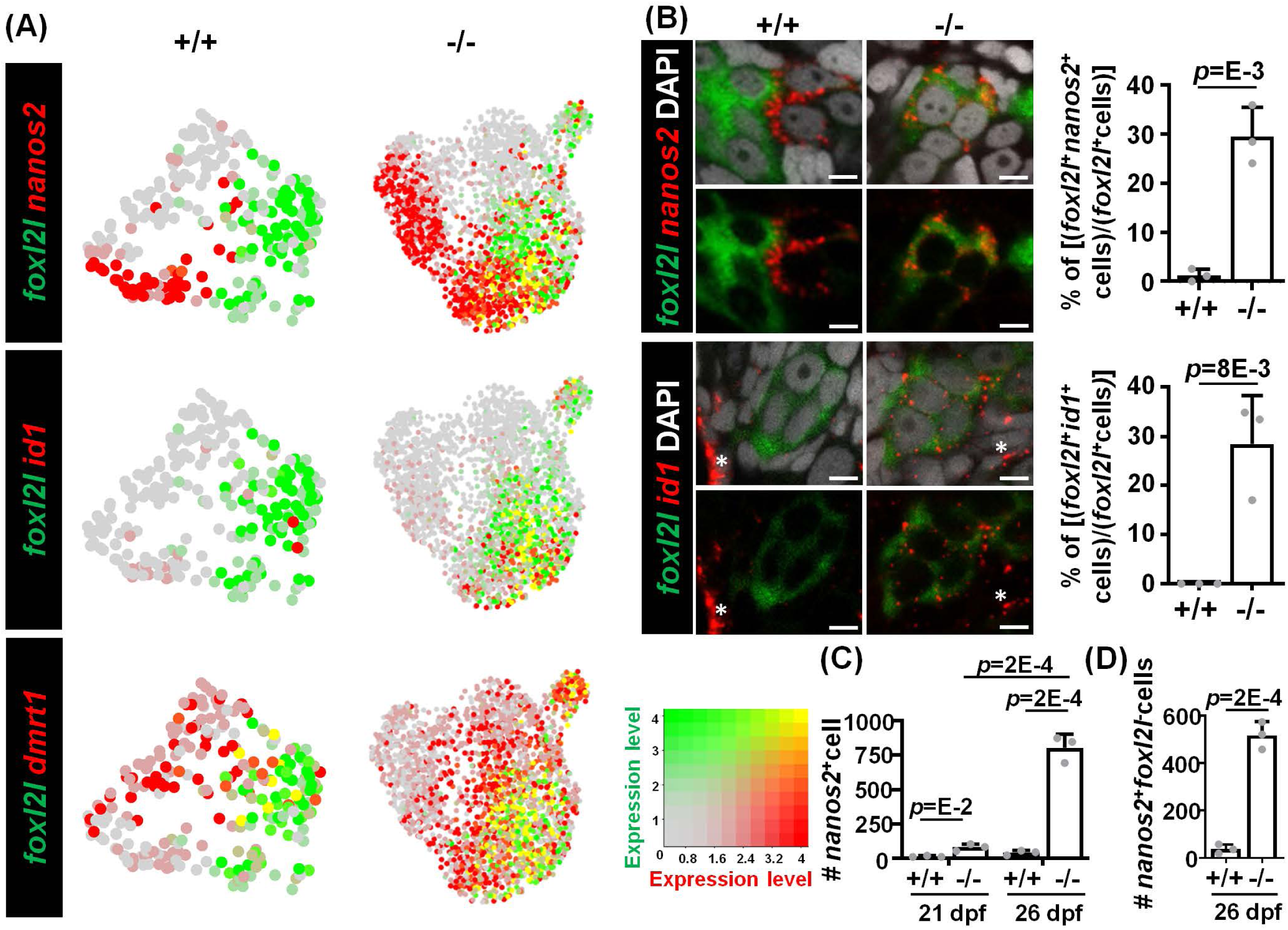
Aberrant co-expression of *foxl2l* with *dmrt1*, *id1* and *nanos2*. **(A)** Visualization of *dmrt1^+^*, *nanos2^+^* or *id1^+^* cells and *foxl2l^+^* cells by UMAP. Left: WT cells in clusters W1 to W5 (+/+). Right: *foxl2l* mutant (-/-) cells. **(B)** Increased proportions of *foxl2l* mutant cells co-expressing *foxl2l* with *nanos2* or *id1*. (Left) Double RNA FISH detecting *nanos2* or *id1* with *foxl2l* in WT (+/+) or *foxl2l^10i/10i^* homozygous mutant (-/-) gonads at 26 dpf. Stained images are shown with (top panels) or without (bottom panels) DAPI in each set. Asterisks represent the expression of *id1* in somatic cell. Scale bars represent 5 µm. (Right) Quantitation of the proportion of double positive cells in each gonad is shown. One dot represents the data from one gonad. **(C)** Increased numbers of *nanos2*-expressing cells in *foxl2l^10i/10i^* homozygous mutant gonad (-/-). One dot represents the data from one gonad. (**D**) Increased number of *nanos2^+^foxl2l^-^*GSC cells in *foxl2l^10i/10i^* homozygous mutant gonad (-/-).

Another gene, *id1*, was barely expressed in WT germ cells, but was abundant in some *foxl2l^+^* mutant germ cells (Figure 5A). Double FISH further detected an increased proportion of *id1^+^foxl2l^+^* cells in the mutant (Figure 5B). Id1 is involved in the maintenance of stemness in various organs (Kantzer et al., 2022; Ying et al., 2003; Zhang et al., 2014), consistent with the hypothesis that mutant Prog-C revert to the GSC property.

Dmrt1 is a male regulator in many vertebrate species (Ge et al., 2017; Matson et al., 2011; Smith et al., 2009). In WT, very few cells co-expressed *foxl2l* and *dmrt1*. However, many mutant cells expressed both *foxl2l* and *dmrt1* (Figure 5A). Similar results can also be observed in scRNAseq data extracted from a public database (Liu et al., 2022)- co-expression of *foxl2l* with *nanos2*, *id1* or *dmrt1* were hardly detected in 40-dpf female germ cells (Supplementary Figure S9).

## Discussion

In this article, we have analyzed development of zebrafish germ cells and elucidated the mechanism controlling sex differentiation in zebrafish by analyzing *foxl2l* mutants. The loss of functional Foxl2l leads to gene dysregulation, a halt of Prog-C development, aberrant reversion to the stem cell status and eventual male development. These results show that *foxl2l* endows Prog-C with female fate and prevent them from reverting to the stem cell fate as depicted in Figure 6.

**Fig. 6.**
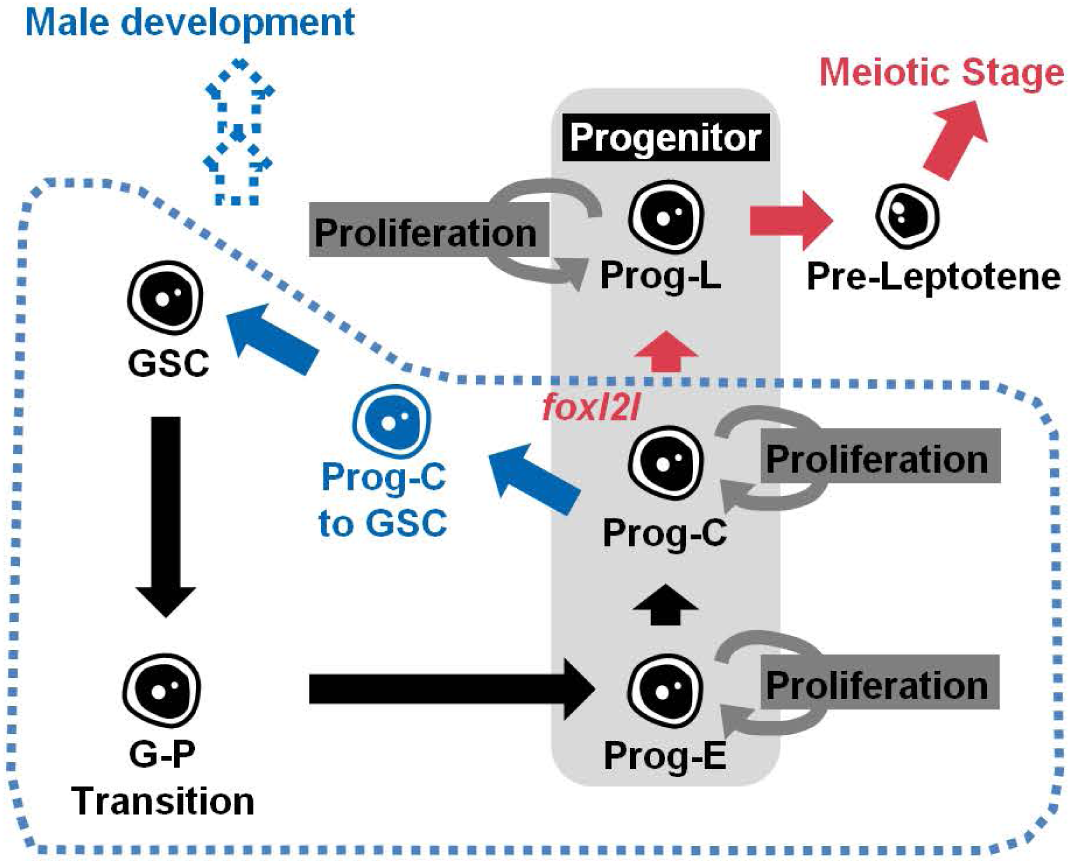
Schematic illustration of the function of Foxl2l during germ cell development in juvenile gonad. During the sex determination period, germ cells develop following the trajectory from germline stem cell (GSC), early progenitor (Prog-E), committed progenitor (Prog-C), late progenitor (Prog-L) to the female meiotic stage. Foxl2l is essential for the maturation of Prog-C and the ensuing Prog-L development. In *foxl2l* mutant, germ cells are arrested at Prog-C. Some mutant Prog-C transit into GSCs. The absence of functional Foxl2l eventually triggers male development. Blue symbols indicate alternative development in *foxl2l* mutants. Red arrows indicate the developmental trajectory of females.

### Developmental trajectory of germ cells

The process of zebrafish germ cell development during the sex differentiation period is poorly understood. Here with scRNAseq analysis, we have depicted the developmental trajectory of germ cells and further characterized GSCs and progenitors. GSCs can be split into two subtypes: *nanos2^hi^cbx7a^lo^*GSC-I and *nanos2^lo^cbx7a^hi^* GSC-II. All GSCs self-renew intermittently to ensure continuous gamete maintenance (Saito et al., 2007). The two subtypes of GSCs may represent two types of daughter cells derived from GSC self-renewal. When GSCs lose stem cell property, they undergo G to P transition for further differentiation into progenitors.

Progenitors include three types: Prog-E, Prog-C, and Prog-L. We show here that Prog-C is the gate toward female determination and will stay at S-phase temporarily before differentiation into Prog-L. In Prog-L, genes such as *rec8a* and *si:ch211-191j22.7* (ortholog of *MEIOSIN*, the meiosis initiator) are highly expressed for the preparation of meiotic entry. Our study delineates the developmental process of zebrafish germ cells during sex differentiation.

### Some zebrafish progenitor germ cells possess female identity

In many vertebrate species, sex determination is induced by sex determining genes expressed in supporting cells (Berta et al., 1990a; Matsuda et al., 2002). The fate of germ cells is determined following the instruction of the supporting cells. In contrast, in zebrafish the germ cell is required for female development, and the depletion of germ cells results in male development (Slanchev et al., 2005). Our study goes beyond this notion and determines that some progenitor germ cells are already sexually differentiated, and Foxl2l in Prog-C dictates female development.

Although Foxl2l is important for female germ cell differentiation in both zebrafish and medaka, there are differences regarding their mutant phenotype between these two fishes. Medaka *foxl2l* (termed *foxl3*) mutant gonads still contain some meiotic germ cells (Nishimura et al., 2015). Zebrafish *foxl2l* deficiency, however, leads to cell arrest at Prog-C as shown in this article. No meiotic cells can be detected in zebrafish *foxl2l* mutant gonads. Furthermore, medaka Foxl2l is involved in meiosis and folliculogenesis (Kikuchi et al., 2020), while zebrafish Foxl2l guides progenitor development towards commitment to the female fate as shown here. It indicates the early establishment of zebrafish germline feminization in the progenitor stage. Zebrafish Foxl2l exerts an earlier role in female differentiation than medaka Foxl2l. The differences between zebrafish and medaka reflect the timing of Foxl2l action between these two species.

### Direct male differentiation from bipotential gonads

Zebrafish testis development has been an unsolved issue. Zebrafish testis can be derived from an ovary-like tissue that undergoes oocyte apoptosis during sex differentiation (Uchida et al., 2002). Here we show that germ cells are arrested in the middle of progenitor development followed by direct male differentiation in *foxl2l* mutant. This indicates that zebrafish testis can be differentiated directly from bipotential gonads without going through the female phase. Our current result is also consistent with the reports showing direct male differentiation (Pan et al., 2022; Tong et al., 2010). Therefore, in addition to female-to-male transition, zebrafish possess another sexual development process similar to most gonocharists whose males and females are derived directly from undifferentiated gonads.

### Foxl2l suppresses genes in male development and stemness

Our scRNAseq analysis has identified *nanos2*, *dmrt1* and *id1*, which may be involved in cell type transition in *foxl2l* mutants. Nanos2 functions in germ cell development in various species (Z. Cao et al., 2019; Kusz et al., 2009; Tsuda et al., 2003). NANOS2 also suppresses meiosis in male mice (Suzuki & Saga, 2008). Therefore, the aberrantly induced *nanos2* in Prog-C of *foxl2l* mutant may participate in the suppression of meiotic genes in addition to the induction of stemness.

Id1, inhibitor of DNA binding 1, is a transcription regulator that lacks DNA binding domain. It can bind basic helix-loop-helix (bHLH) proteins to prevent their binding to DNA (Benezra et al., 1990). Id1 is essential for homeostasis and the maintenance of stem cells in several cell types (Hong et al., 2011; Kantzer et al., 2022; Ying et al., 2003; Zhang et al., 2014). Although the function of Id1 in the testis remains unclear, its paralog Id4 regulates self-renewal of spermatogonial stem cells in the testis (Sablitzky et al., 1998; Wang et al., 2018). Thus, the enhanced *id1* expression in the mutant Prog-C germ cells may inhibit genes controlling differentiation and guide cells toward stem cell fate in the *foxl2l* mutant.

Dmrt1 regulates male differentiation in many animals. In Nile tilapia (*Oreochromis niloticus*), Dmrt1 and Foxl2l antagonize each other to determine germline sex (Dai et al., 2021). Zebrafish appears to behave similarly. As a male-promoting gene (Webster et al., 2017), zebrafish *dmrt1* is aberrantly upregulated in *foxl2l* mutants as shown here. This aberrant *dmrt1* expression may direct *foxl2l* mutants toward male development.

We show here that Foxl2l may ensure female differentiation by preventing stemness and antagonizing male development. It does so by suppressing stem-cell genes (*id1* and *nanos2*) and male gene *dmrt1*. In the absence of functional Foxl2l, the upregulation of *id1* and *nanos2* triggers progenitor transition to GSC while the upregulation of *dmrt1* induces male differentiation.

### The alternative development of *foxl2l* mutant germ cell

We have detected aberrantly enriched *tp53* and *zgc:158463* in distinct *foxl2l* mutant cells in I3 or I6. Tumor suppressor protein p53, encoded by *TP53*, induces cell cycle arrest and apoptosis. Furthermore, p53 also regulates the differentiation of airway epithelial progenitors (McConnell et al., 2016), embryonic stem cells (Lin et al., 2005), and brown adipocytes (Molchadsky et al., 2013). Thus, p53 may assist alternative differentiation of mutant germ cells. We have also identified an uncharacterized gene, *zgc:158463*, which is specifically expressed in *foxl2l* mutant cells in I3 and I6. Its high expression suggests that it may be involved in the male differentiation of mutant cells.

## Materials and Methods

### Animals

Wildtype zebrafish *Danio rerio*, TL and Nadia strains, *piwil1:egfp* transgenic (Pan et al., 2022) and *foxl2l^-/-^* mutant fish were maintained in system water (28.5°C, pH7.2, conductivity 500 µS/cm) according to the standard protocol (Westerfield, 2000). For embryos, fertilized eggs were raised in 100-mm petri dish with 100 eggs per dish until 4 dpf (days post fertilization). Larvae were then transferred to 1-liter beaker with 60 larvae in 600 ml system water and fed with paramecia until 12 dpf. At 8 dpf, larvae density was reduced to a half and fed with brine shrimps twice daily. At 21 dpf, larvae were transferred to system tank. For sample collection, the growth of larva was determined according to both age and body length (Nüsslein-Volhard, 2002). Zebrafish were handled under the guidelines of Institutional Animal Care and Utilization Committee of Academia Sinica and National Laboratory Animal Center of National Applied Research Laboratories.

### Generation of *foxl2l* mutant

The gRNA sequences targeting the forkhead domain of *foxl2l* were designed using CHOPCHOP (https://chopchop.cbu.uib.no) and CRISPRscan (https://www.crisprscan.org). Target sites with high efficiency, low off-target effect and the sequence of NGG at the 3’ end were chosen. To get higher knockout efficiency, two gRNA with 50 pg each and 200pg of TrueCut Cas9 Protein v2 (Life Technologies, Cat. No. A36498) were co-injected into embryo at two-cell stage. F0 fish were crossed with WT fish to generate stable mutant lines. To determine the genotype, the fin of F1 offspring was heated in 50 mM NaOH at 95°C followed by the addition of 1/10 volume of 1 M Tris-HCl pH 8 on ice for 5 mins before DNA amplification by PCR (F, 5’- AGTAAACCTGAAGCACACCTGG-3’, R, 5’-CATCCCTTTTTGTTCTTCTCGT-3’) using Fast-RunTM 2x Taq Master Mix without Dye (Protech Technology Enterprise CO., Cat. No. PT-TMM228-D). The size difference between WT and mutant alleles was determined from capillary electrophoresis, run by QIAxcel DNA Screening Kit (QIAGEN, Cat. No. 929004), detected by High Performance Nucleic Acid Analyzer (eGENE HDA-GT12) and analyzed by QIAxcel BioCalculator software.

### RNA analysis and Plasmid

Tissue RNA was extracted by TRIZOL reagent (Invitrogen, Cat. No. 15596026). cDNA was reverse transcribed by Maxima First Strand cDNA Synthesis kit (Fermentas Int, Cat. No. K1641) from 1 µg of RNA.

For probe synthesis, templates were either constructed into vector (for *foxl2l*, *nanos2* and *id1*) or amplified by PCR (for *rec8a*, *obf3b* and *tekt1*). For construction, cDNA form ovary (for *foxl2l* and *nanos2* probe) or 24-hpf larvae (hours post fertilization) (for *id1*)(Addgene) were amplified by PCR (*foxl2l*: F, 5’-CTTTCCACCTGTACCGTGCG-3’, R, 5’-CAGTCAGCACCGAGGTTTGC-3’; *nanos2*: F, 5’-GACGGATCCCATGGGCAAAACACACCTAAAACA-3’, R, *id1*: F, 5’-TGGTGAACTGTCATCGCACT-3’, R, 5’-AGCGTTCACATCATATGGCA-3’) using Taq DNA polymerase (Roche, Cat. No. 11146165001). Fragments of *foxl2l* and *id1* were cloned into *pGEMT-easy* plasmid by TA cloning while *nanos2* fragment was cloned into pCS2^+^ plasmid by *BamHI* and *KpnI*. Plasmid was cut and antisense probe was synthesized by in vitro transcription with Dig RNA Labeling Mix (Roche, Cat. No. 11277073910) or Fluorescein RNA labeling mix (Roche, Cat. No. 11685619910) together with T7 RNA polymerase (Roche, Cat. No. 10881775001), SP6 RNA polymerase (Roche, Cat. No. 10810274001) or T3 RNA polymerase (Roche, Cat. No. 11031163001)(*foxl2l*: *NcoI*, SP6; *nanos2*: *BamHI*, T3; *id1*: *SacII*, SP6). For *rec8a, obf3b* and *tekt1*, cDNA containing T7 promoter were amplified by PCR (*rec8a*: F, GAGTATTTAGGTGACACTATAGACAATTCCCCCTCAGCAACC, R, GAGTAATACGACTCACTATAGGGGATGCACCGGTGATTTGTGC; *obf3b*: F, 5’-GAGTATTTAGGTGACACTATAGGGGGCAACTGGAATGAATAA-3’, R, 5’-GAGTAATACGACTCACTATAGGGACTACGACCGCTGAAGGAGA-3’; *tekt1*: F, 5’-GAGTATTTAGGTGACACTATAGGGAGGATCCAGGACATCAAA-3’, R, 5’-GAGTAATACGACTCACTATAGGGCCTTCTCGGCTTTGCTAATG-3’) using Fast-RunTM 2x Taq Master Mix without Dye (Protech Technology Enterprise CO., Cat. No. PT-TMM228-D). After purification by QIAquick PCR Purification Kit (QIAGEN, Cat. No. 28106), antisense probe was synthesized by in vitro transcription as described above.

For the generation of *foxl2l* mutant lines, the *pT7-gRNA*-*foxl2l#1* and *pT7-gRNA*-*foxl2l#2* plasmids containing oligo against two different target site of *foxl2l* were constructed. After annealing oligos pairs (*foxl2l#1*: F, 5’-TAGGAGCTGGATGAATGAAACG-3’; R, 5’-AAACCGTTTCATTCATCCAGCT-3’; *foxl2l#2*: F, 5’-TAGGAGCCACGTACGAATAAGG-3’, R, 5’-AAACCCTTATTCGTACGTGGCT-3’) *foxl2l#1* and *foxl2l#2* fragments were cloned into *pT7-gRNA* (Addgene) through one-step digestion and ligation respectively. Reagents (400ng of *pT7-gRNA*, 0.25 µM annealed oligo, 1x NEBuffer 3.1 (New England Biolabs, Cat. No. B7203), 1x T4 DNA ligase buffer (Promega Co., Cat. No. C126B), 1.5 U T4 DNA ligase (Promega Co., Cat. No. M180A), 5 U *BsmBI* (New England Biolabs, Cat. No. R0580), 3 U *BglII* (New England Biolabs, Cat. No. R0144S) and 6 U *SalI* (New England Biolabs, Cat. No. R0138S)) were incubated with the following condition, 20 mins in 37°C and 15 mins in 16°C for three cycles followed by 10 mins in 37°C, 15 mins in 55°C, 15 mins in 80°C and cooling in 4°C, for plasmid construction. Then the plasmids were linearized by *BamHI*, and gRNA was synthesized by MegaShortscript T7 transcription Kit (Thermo Fisher Scientific, Cat. No. AM1354).

### RNA *in situ* hybridization

Whole mount *in situ* hybridization is performed using a published protocol (Thisse & Thisse, 2008). Larvae or tissues were fixed in 4% paraformaldehyde (PFA) overnight at 4°C before the removal of head, tail and intestine. The dissected samples went through dehydration, rehydration, permeation, hybridization, antibody incubation and staining. For permeation, larvae were treated with 10 µg/ml Proteinase K (Roche, Cat. No. 03115887001) for 10 mins while adult tissues were treated for 40 mins. For probe hybridization, samples were incubated with 1 ng/µl digoxigenin (DIG)-labeled probe at 70°C overnight. For color staining *in situ* hybridization, the antibody incubation was performed with 1:5000 diluted Anti-Dig-AP antibody (Roche, Cat. No. 11093274910) at 4°C overnight. Samples were stained with BM-purple (Roche, Cat. No. 11442074001). Fluorescent *in situ* hybridization was performed as described (Brend & Holley, 2009) with 555 Styramide Kit (AAT Bioquest Inc., Cat. No. 45027) and 488 Tyramide Conjugate (Biotium Inc, Cat. No. 92171) followed by DAPI staining with 1:20000 dilution. For photography, samples were mounted in 85% glycerol.

### Photography

Photography was performed using AxioImager Z1 upright microscope (Carl Zeiss Inc.) with AxioCam HRc digital camera. Image was processed by Axiovision 4.7 software (Carl Zeiss Inc.). For fluorescent samples, photography was performed by LSM 780 Confocal Microscope (Carl Zeiss Inc.).

### Mating test

Adult female with different genotypes were mated with WT males individually from 4 months of age. Ten matings were performed with 7- to 14-day intervals. To avoid individual difference, the first three matings were considered as practice and were not included in the experimental counting. Fertilization rate was calculated at 24 hours post fertilization (Fertilization rate: (number of fertilized egg)/(total number of egg)). Each experimental group contained three to seven mating pairs.

### Histology

Larvae, ovary and testis were fixed in Bouin’s solution at 4°C overnight before 6 washes in PBS for 30 mins each, serial dehydration from 50% to 100% ethanol and incubation with xylene and paraffin twice each. Then the samples were embedded in paraffin before being sectioned with 6-µm thickness. The sectioned samples were stained with hematoxylin and eosin for histological observation.

### Immunofluorescence

Larvae were fixed in 4% PFA overnight before removal of head, tail and intestine. The dissected samples went through three washes of PBS with 0.2% Triton X-100 (PBST) for 5 mins each, blocking with normal goat serum for 1 hour at room temperature, incubating with primary antibody (Vasa (GeneTEX, Cat. No. GTX128306), 1:200 dilution; Sycp3 (Abcam, Cat. No. ab150292), 1:100 dilution) 4°C overnight, three washes in PBST for 30 mins each and incubating with 1:200 diluted secondary antibody (Alexa Fluor 546 Donkey anti-Rabbit IgG (Invitrogen, Cat. No. A-10040); Alexa Fluor 488 Goat anti-Rabbit IgG (Invitrogen, Cat. No. A-11034)) together with 1:20000 diluted DAPI. Finally, the samples were mounted in 85% glycerol for photography.

### Transcriptome analysis of single cell RNA-seq

Germ cells from 26-dpf WT or *foxl2l*^10i/10i^ zebrafish in *piwi1:EGFP* transgene background were collected for single cell RNA-seq. To isolate EGFP^+^ germ cell, the trunk of 20 fish were collected and dissociated in 0.4% collagenase (Worthington Biochemical, Cat. No. LS004188) and 0.5% trypsin (Life Technologies, Cat. No. 15400-054) in 1 mL phosphate buffer saline (PBS) at 28°C for 60 mins. After dissociation, cells were supplemented with fetal bovine serum (FBS) (Life Technologies, Cat. No. 10437028), centrifuged at 600g, 4 min, and the pellet was suspended in FACS™ Pre-Sort Buffer (BD Biosciences, Cat. No. 563503) before being filtered through cell strainers. 1mg / 1 μL of Propidium iodide (Sigma-Aldrich, Cat. No. P4170) was added to the eluate to stain dead cells, and around 75% survival rate was obtained in both genotypes. EGFP^+^ cells were collected into 60% Leiboviz’s L-15 medium (Thermo Fisher Scientific, Cat. No. 11415064) by FACSAria IIIucell sorter (BD Biosciences) and supplemented with 10% FBS. Total 4781 cells for *foxl2l*^+/+^ and 8673 cells for *foxl2l*^10i/10i^ samples were obtained upon counting by a cell sorter. After centrifugation at 300 g for 5 min and resuspension in PBS with 0.04% bovine serum albumin (Sigma-Aldrich), the recovered cell number was 3231 in *foxl2l*^+/+^ and 4019 in *foxl2l*^10i/10i^. Finally, the single cell cDNA libraries were constructed by Chromium Next GEM Single Cell 3’ Library Construction Kit v3.1 kit (10x Genomics) following the manufacturer’s protocols with 12 cycles for cDNA amplification and 14 cycles for library amplification.

Sequencing was performed by NextSeq 500 (Illumina) with 150 cycles of 28 x 91 paired-end reads. The raw base call (BCL) files were transferred to FASTQ file by the cellranger mkfastq pipeline from Cell Ranger (10x Genomics). The reads were mapped to zebrafish genome GRCz11 and gene expression was obtained by cellranger count from Cell Ranger (10x Genomics).

### Statistical analysis

All the quantitative data presented as mean mean±SD. The data was analyzed by Student’s *t*-test using GraphPad Prism 8.

### Single-cell transcriptome analysis

#### Count matrix generation

Raw count matrices were generated by Cell Ranger v3.1.0 (Zheng et al., 2017) using the Zebrafish reference genome GRCz11 and reference transcriptome ensemble release v101. The count matrices were then stored as Seurat objects v3.2.3 (Stuart et al., 2019).

#### Quality control

A two-step quality control approach was conducted individually for each sample. First, we conducted a quality control at the cell level. Cells with UMI counts < 10,000 or mitochondrial gene expression percentages 2: 10% were marked as low-quality cells. Multiplet detection was performed on the remaining cells using the R package *scds* (Bais & Kostka, 2020). Cells with a hybrid score 2: the outer fence were marked as multiplets. Second, we performed quality control at the cluster level to screen out the potential low-quality cells that passed the hard thresholds at the cell level. A low-resolution clustering was performed using Seurat v3.2.3. Clusters containing more than 80% low-quality cells or multiplets were marked as low-quality clusters.

Together, cells that were marked as low quality, multiplets, and cells in low-quality clusters were excluded from the subsequent analysis.

#### Data preprocessing and cell cycle scoring

The count matrices of the remaining 769 cells from the WT and 2,399 cells from the mutant were merged and normalized using the R package *scran* (Lun et al., 2016). After normalization, gene expression data of cells from WT and mutant samples were organized into individual Seurat objects. The function *CellCycleScoring* in Seurat v3.2.3 was used for cell-cycle scoring (Supplementary Figure S10). Briefly, all genes were first sorted by their mean expression and split into bins. For each S- and G2M-specific gene, relative expression was calculated by comparison with other genes in the same bin. The S-score and G2/M-score were calculated as the average of the relative expression of all genes in the lists of S- and G2M-specific genes.

#### Cell clustering analysis of WT cells

We used the *FindVariableFeatures* in Seurat v3.2.3 to identify the top 2,500 highly variable genes in each sample. The principal components of these highly variable genes on scaled expression matrices were derived after their cell-cycle scores were regressed out (Supplementary Figure S11). We used the leading 30 principal components for UMAP embedding. The Leiden algorithm was applied to identify cell clusters. The optimal resolution, determined by the highest average silhouette score, was chosen from a range of feasible values.

#### Trajectory and pseudo-time analysis of WT cells

The data matrix, along with the cell clusters and UMAP embedding, were converted to the CDS format and subsequently analyzed using Monocle 3 (J. Cao et al., 2019) to obtain the trajectory that encompasses all clusters. We then used the germline stem cell (GSC) marker *nanos2* to identify the root and the order of the trajectory and to estimate the pseudo-time for each WT cell.

#### Cluster marker identification

We used the *top_markers* in Monocle 3 (J. Cao et al., 2019) to identify the cluster markers whose high expression discriminates cells in the target cluster from the other cells with a high degree of specificity. For each cluster, genes that satisfy the following criteria: (i) specificity ≥ 0.2, (ii) Q-value < 0.01, and (iii) having non-zero counts in ≥ 80% cells in the cluster were selected as markers. Top marker genes were selected based on the highest specificities within WT clusters.

#### Assigning the developmental stages to mutant cells

The function *TransferData* in Seurat v3.2.3 was utilized to transfer the developmental stage from the WT cells to the mutant cells. Initially, principal components (PCs) were derived from the WT cells based on the expression of their cluster markers. Mutant cells were then projected to the same PCs. The PCs and the projected PCs were used to identify the mutual nearest neighbors across mutant cells and WT cells. These mutual nearest neighbors were considered anchors and scored based on the consistency across the neighborhood structure of each dataset. Lastly, the developmental stage of each mutant cell was determined using a weighted voting mechanism. This process considers the stages of WT cells in every anchor, with votes weighted according to the anchor score and the proximity between the mutant cell and its corresponding anchor mutant cell.

#### Co-clustering analysis of mutant cells and WT cells from GSC to Prog-C

To integrate mutant cells and WT cells from GSC to Prog-C, the function *DataIntegration* in Seurat v3.2.3 was applied. Briefly, both datasets were projected to a shared space by performing canonical correlation analysis. Mutual nearest neighbors were then identified as anchors, representing pairs of mutant and WT cells sharing a similar expression pattern. Anchors were scored based on the consistency across the neighborhood structure of each dataset. For each WT cell, a correction vector was created based on the proximity to every anchor and the anchor score. The WT expression matrix was corrected accordingly and integrated with that of mutant cells for the same clustering procedure as previously described. Mutant-specific marker genes were identified using the top_markers function with the following criteria: (i) specificity ≥ 0.25, (ii) Q-value < 0.01, (iii) presence in ≥ 80% mutant cells for each integrated group, and (iv) not included in the WT cluster marker gene list.

#### Composition analysis

Because each stage has its own overall rate of prevalence in the entire cell population, we adjusted for the unevenness of stage prevalence in determining how dominant a stage is in an integrative group. Fisher exact test was used to find out which integrative groups were enriched with what stages of cells. Dominant stages for an integrative group are defined as those with significantly higher proportions of cells than their prevalence rates. Specifically, we calculated the proportions of developmental stages for both WT and mutant cells in each group and then conducted Fisher’s exact tests for each stage within each group. The resulting *P*-values were adjusted by the Benjamin-Hochberg method for controlling for false discovery rates, which were set at 0.05. The stages with significant enrichment are referred as the dominant stages after prevalence rate calibration, or dominant stages for short.

#### Differential expression analysis

To identify DEGs during development, we compared gene expression levels between mutant and WT cells at the same dominant stage within each integrated group. Other cells not included in the comparison are shown in gray on UMAP (Figure 4A). The Wilcoxon rank sum test was performed using the function *FindMarkers* in Seurat v3.2.3. Genes that satisfy the following criteria: (i) |fold-change| ≥ log2(2.5), (ii) Q-value < 0.05, and (iii) gene expression detected in ≥ 50% of cells in either WT or mutant, were selected as DEGs. The DEGs in each stage were ordered by the fold change.

#### Reprocessing of the public scRNA-seq dataset from Liu et. al

The processed data were retrieved from the designated repository. To visualize the expression of GSCs and progenitors of WT cells in 40 dpf, we selected a subset of cells with sufficient UMI counts (2: 7,000) from clusters 0 and 2 and lacked *zp3.2* gene expression. After regressing out the cell-cycle scores, we used the leading 30 principal components for UMAP embedding.

#### Data availability

Analysis code and data for single-cell transcriptomic data analysis are available with NCBI GEO (https://www.ncbi.nlm.nih.gov/geo/query/acc.cgi?acc=GSE173718).

## Acknowledgments

We would like to thank the help of Flow Cytometry Core (Institute of Biomedical Sciences), Genomics Core and Bioinformatics Core (Institute of Molecular Biology Academia Sinica) in scRNAseq study, and Taiwan Zebrafish Core Facility (NSTC 112-2740-B-400-001), Imaging Core (Institute of Molecular Biology Academia Sinica) for confocal imaging.

## Author Contribution

C.-w.H., H.H, C.-H. Y., Y.-w. W., K.-C. L., B.-c.C. designed the experiments, analyzed data, and wrote the manuscript jointly. Y.-w. W. generated knockout fish. C.-H. Y. dissected germ cells for scRNAseq. H.H. and K.-C. L. performed bioinformatic analysis. C.-w.H performed staining to validate the scRNAseq result. B.-c.C and K.-C. L. secured funding and oversaw the execution of the project.

## Funding

This work was funded by grants from National Health Research Institutes (NHRI-EX107-10506SI) and Ministry of Science and Technology (MOST 108-2311-B-001 -038 -MY3 to BcC).

## Supporting Information

The supporting information includes supplementary materials and methods, supplementary references, supplementary figures and legends.

---

**Supplementary Fig. S1.**
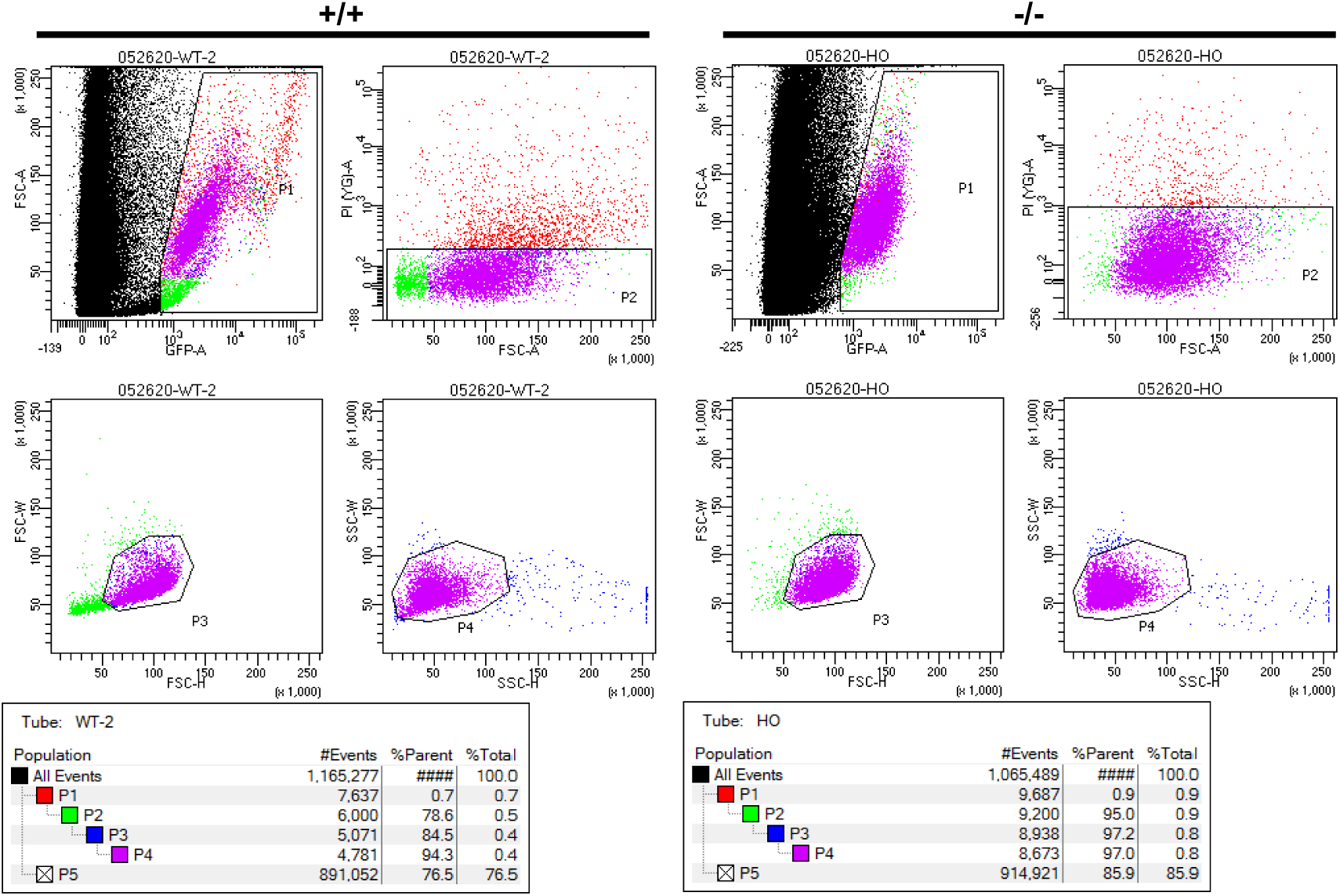
Germ cell sorted by Fluorescence-Activated Cell Sorting (FACS). Sorting of fluorescent germ cell from 26-dpf gonad by FACS. Cells within the rectangular or irregular box were selected. Pseudocolors indicates four different gating, P1 to P4. After gating, germ cells from WT (+/+) or *foxl2l^10i/10i^* (-/-) with *piwil1:egfp* transgenic background were collected.

**Supplementary Fig. S2.**
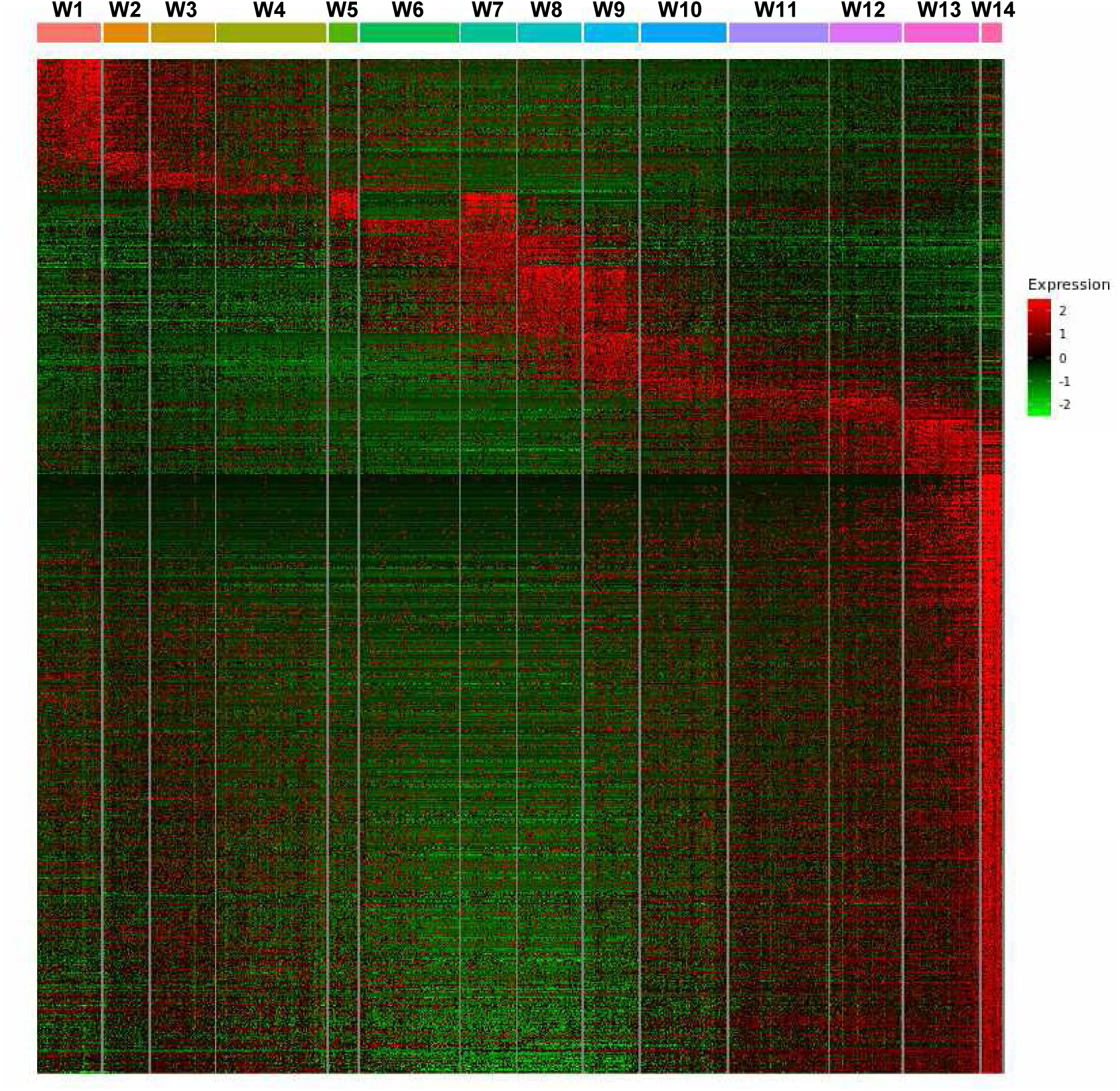
Analysis of WT clusters. Expression of marker genes (Y-axis) in each WT cluster (X-axis) in a heatmap. Genes that reach the criteria (fraction≥ 0.8, specificity ≥ 0.2, *q-*value ≤ 0.01) are defined as marker genes. Expression value is visualized with range between -2 to 2.

**Supplementary Fig. S3.**
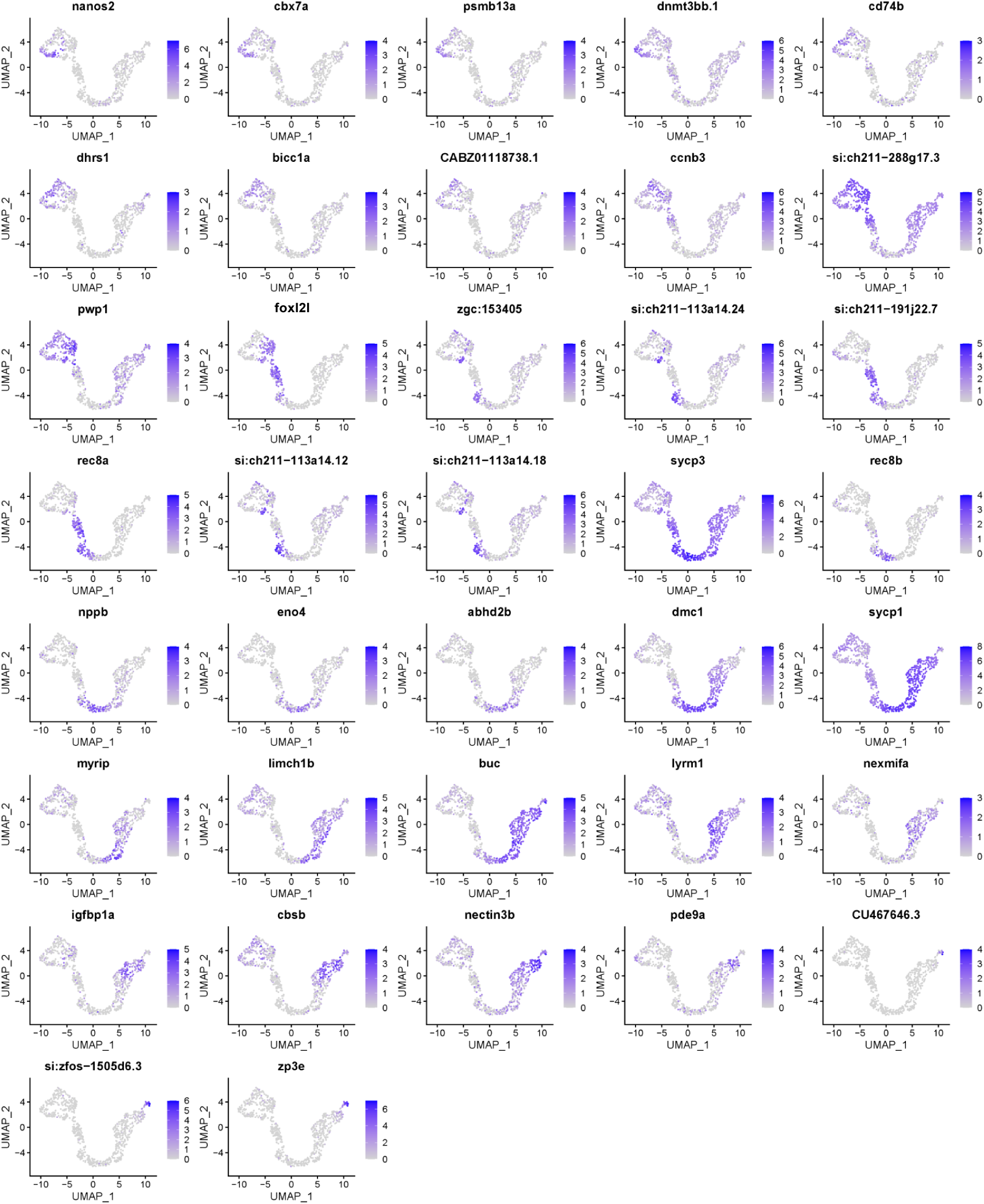
Expression profiles of top and known marker genes in WT germ cell at 26 dpf. Expression of representative WT marker genes shown by Uniform Manifold Approximation and Projection (UMAP). The scales of expression levels are shown at the right of each UMAP.

**Supplementary Fig. S4.**
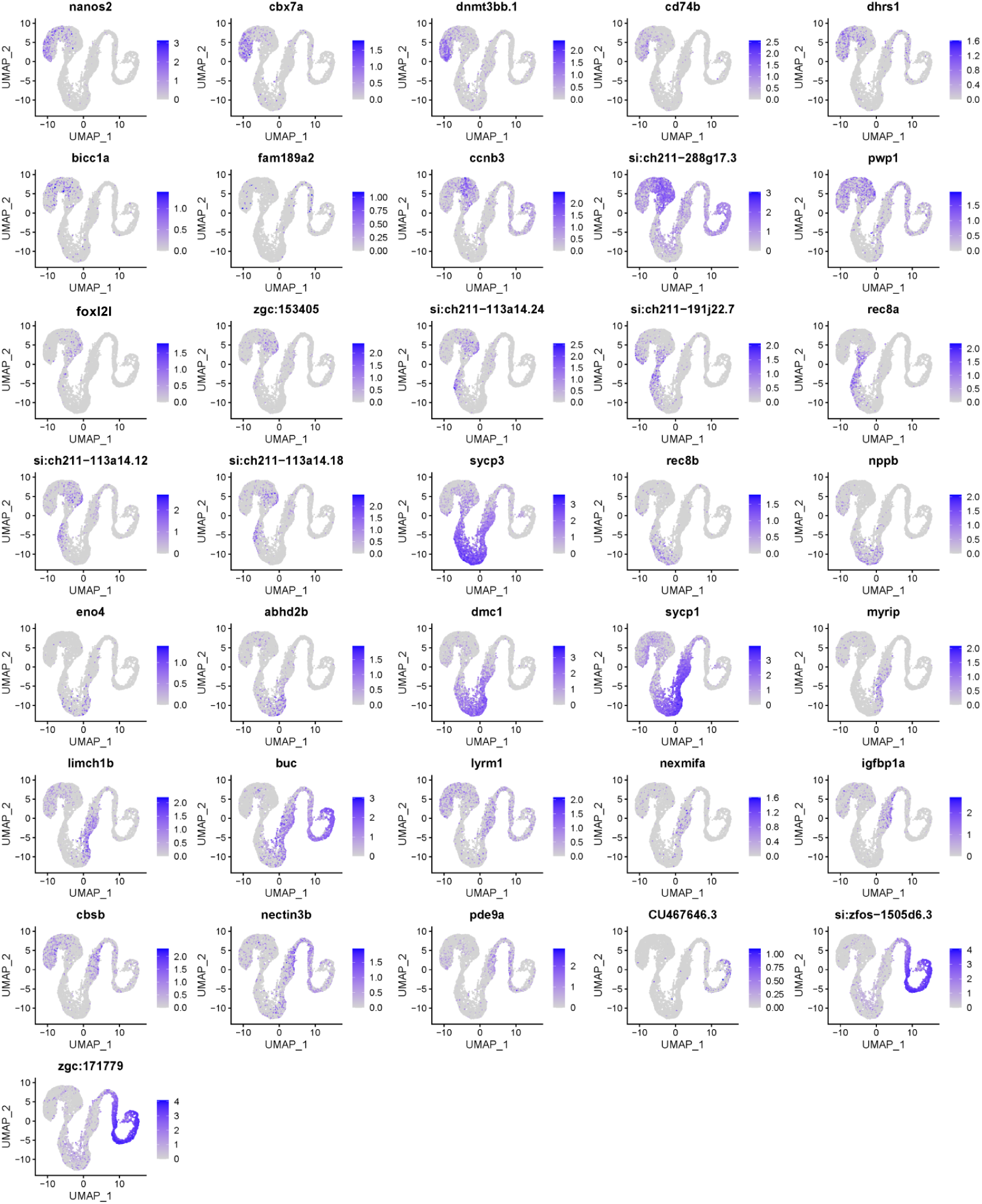
Expression profiles of top and known WT markers extracted from the database. Expression of representative WT marker genes in 40-dpf germ cell obtained from Y Liu, et al. (Liu et al., 2022) shown by UMAP. The representative WT markers were selected from our 26-dpf scRNAseq analysis. The scales of expression levels are shown at the right of each UMAP.

**Supplementary Fig. S5.**
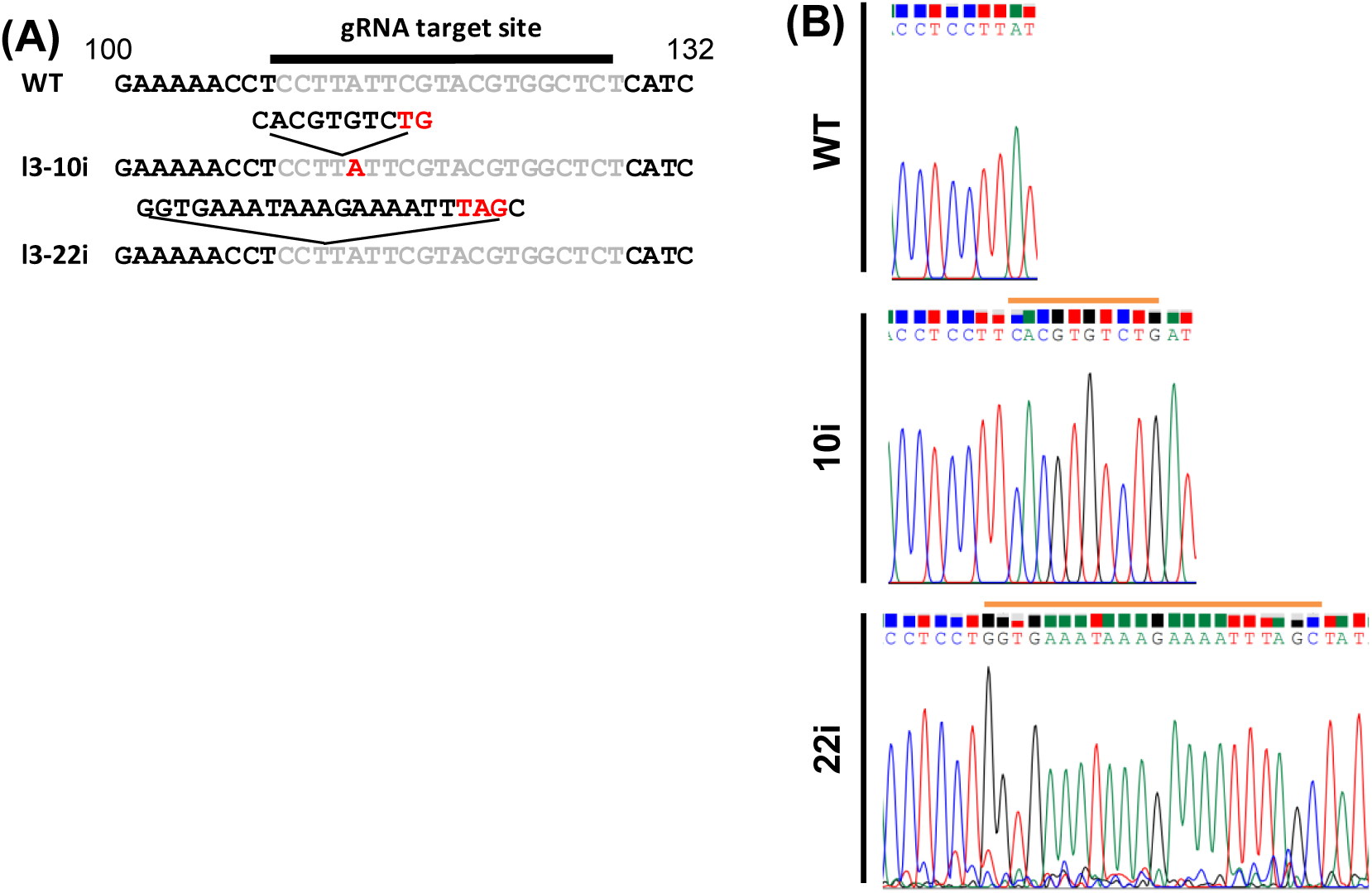
Generation and the phenotype analysis of *foxl2l* mutant. **(A)** The *foxl2l* sequence of WT and two *foxl2l* mutant lines, 10 base pairs insertion (10i) and 22 base pairs insertion (22i). Gray letters represent gRNA targeting region. Red letters represent stop codons. **(B)** Sequencing results of WT and two *foxl2l* mutant lines, 10i and 22i. The inserted sequences are marked with horizontal orange lines.

**Supplementary Fig. S6.**
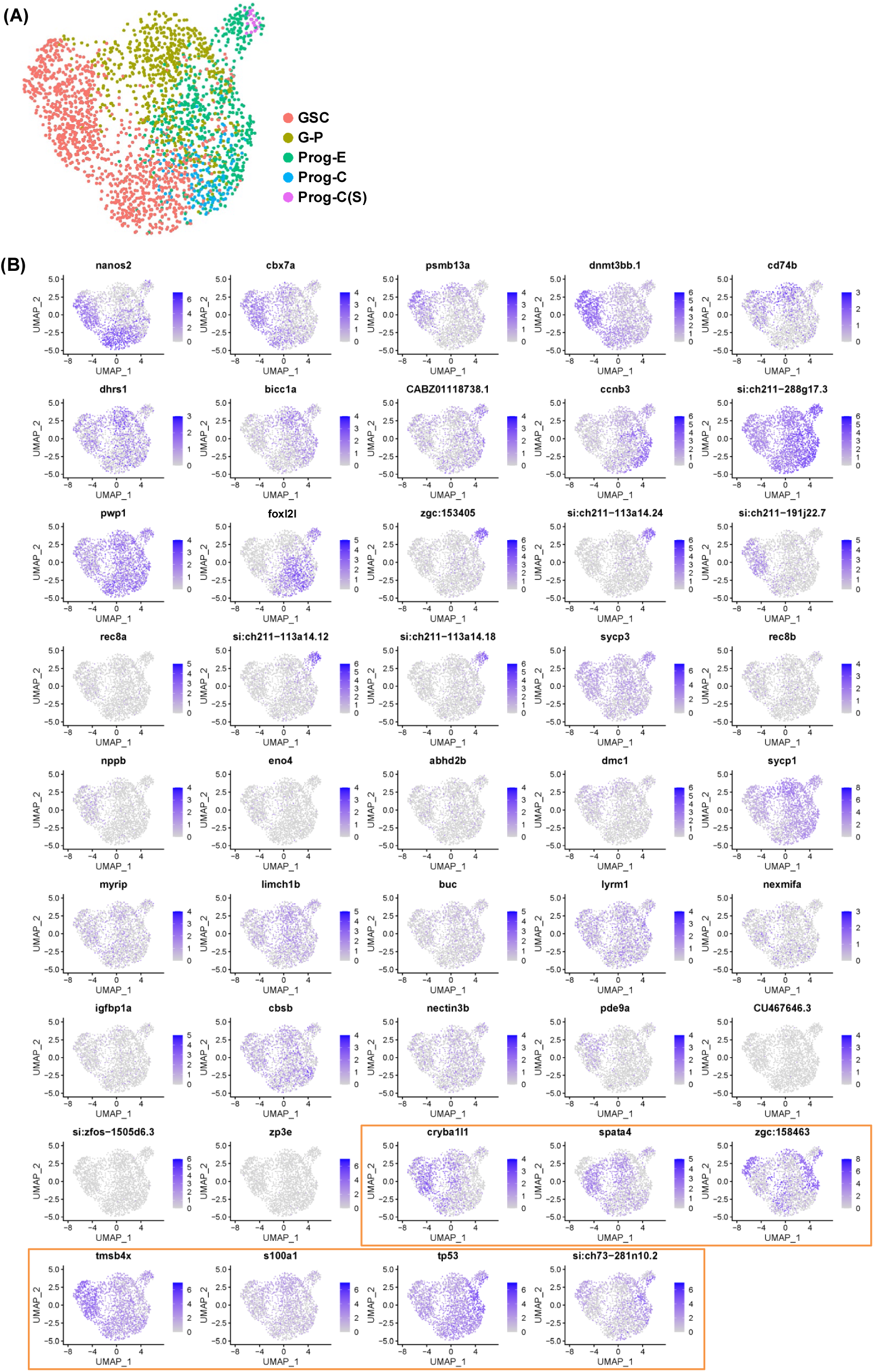
Transcriptome analysis of mutant germ cells. **(A)** Mutant cells shown in UMAP. Color of each mutant cell represents annotated developmental stage. **(B)** Expression of top marker genes for WT and 26-dpf mutant (in orange box) shown by UMAP. The scales of the scores are shown at the right.

**Supplementary Fig. S7.**
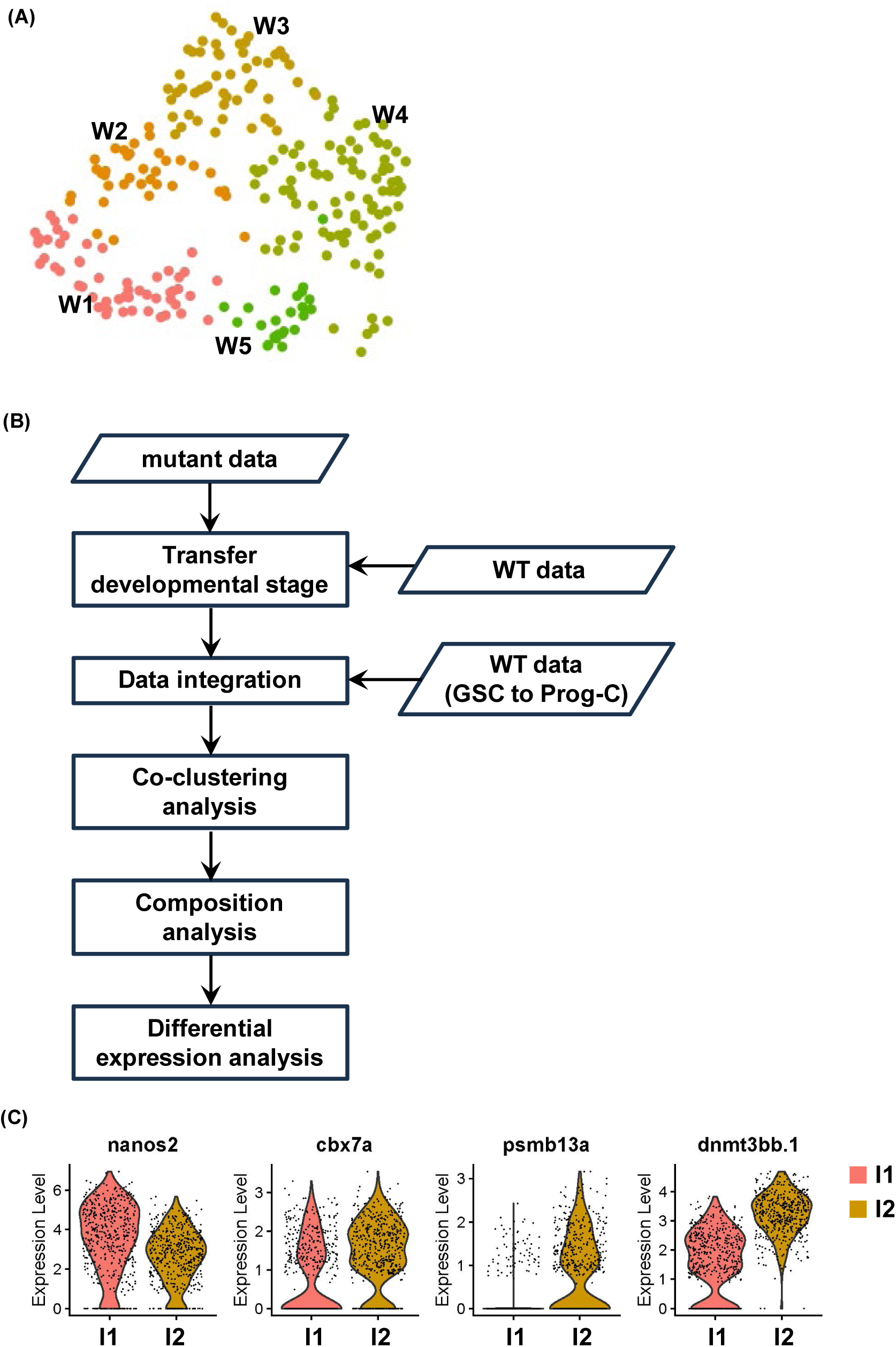
Transcriptome analysis of integrated groups. **(A)**WT germ cells in cluster W1-W5 visualized by UMAP. **(B)** The flowchart of the co-clustering analysis. **(C)** Split-violin plots showing the distribution of cells expressing four top GSC markers in integrated groups I1 and I2.

**Supplementary Fig. S8.**
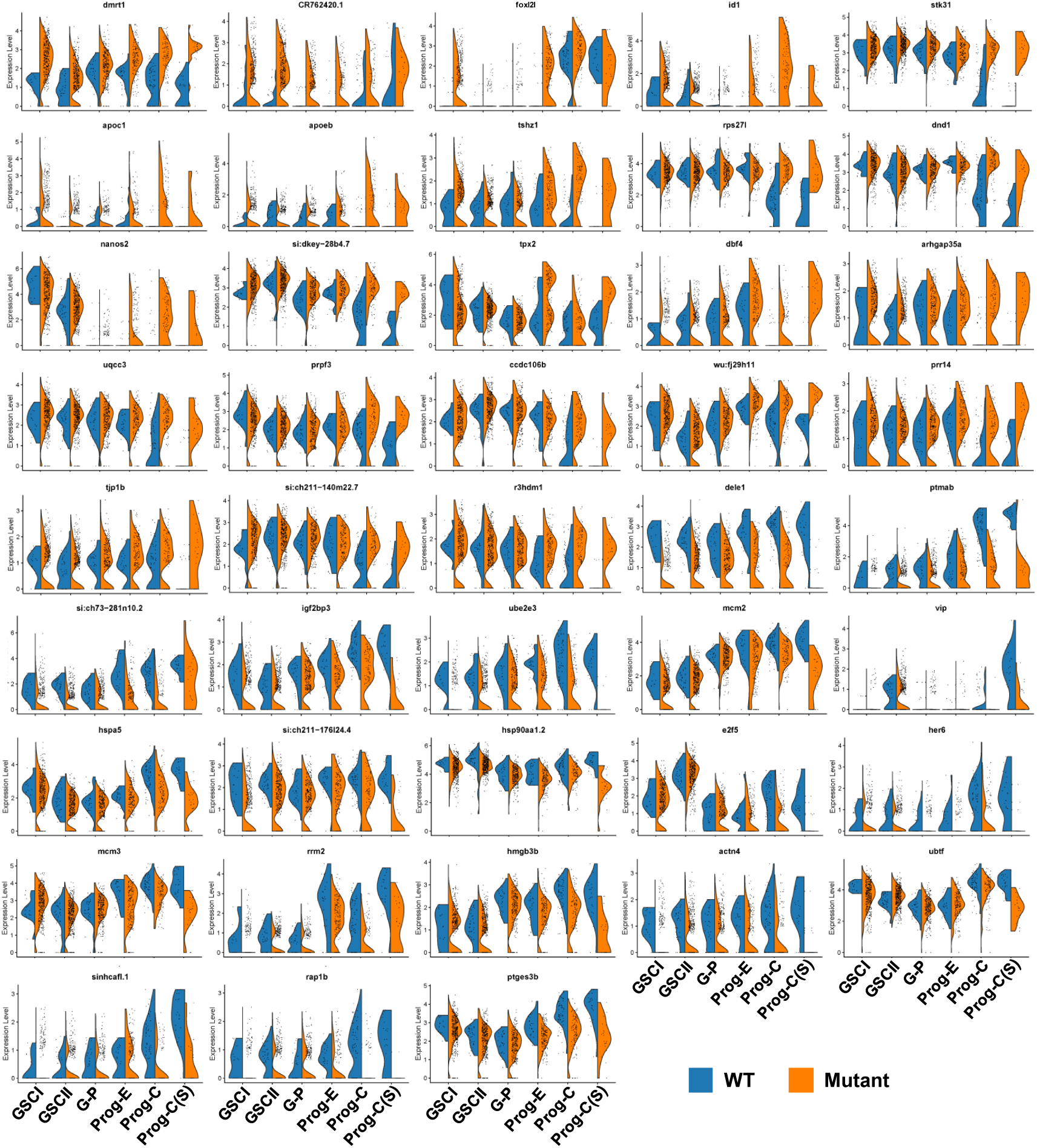
Expression of DEGs in WT and mutant germ cell. Split-violin plots showing the distribution of WT and mutant cells expressing DEGs at different developmental stages (X-axis).

**Supplementary Fig. S9.**
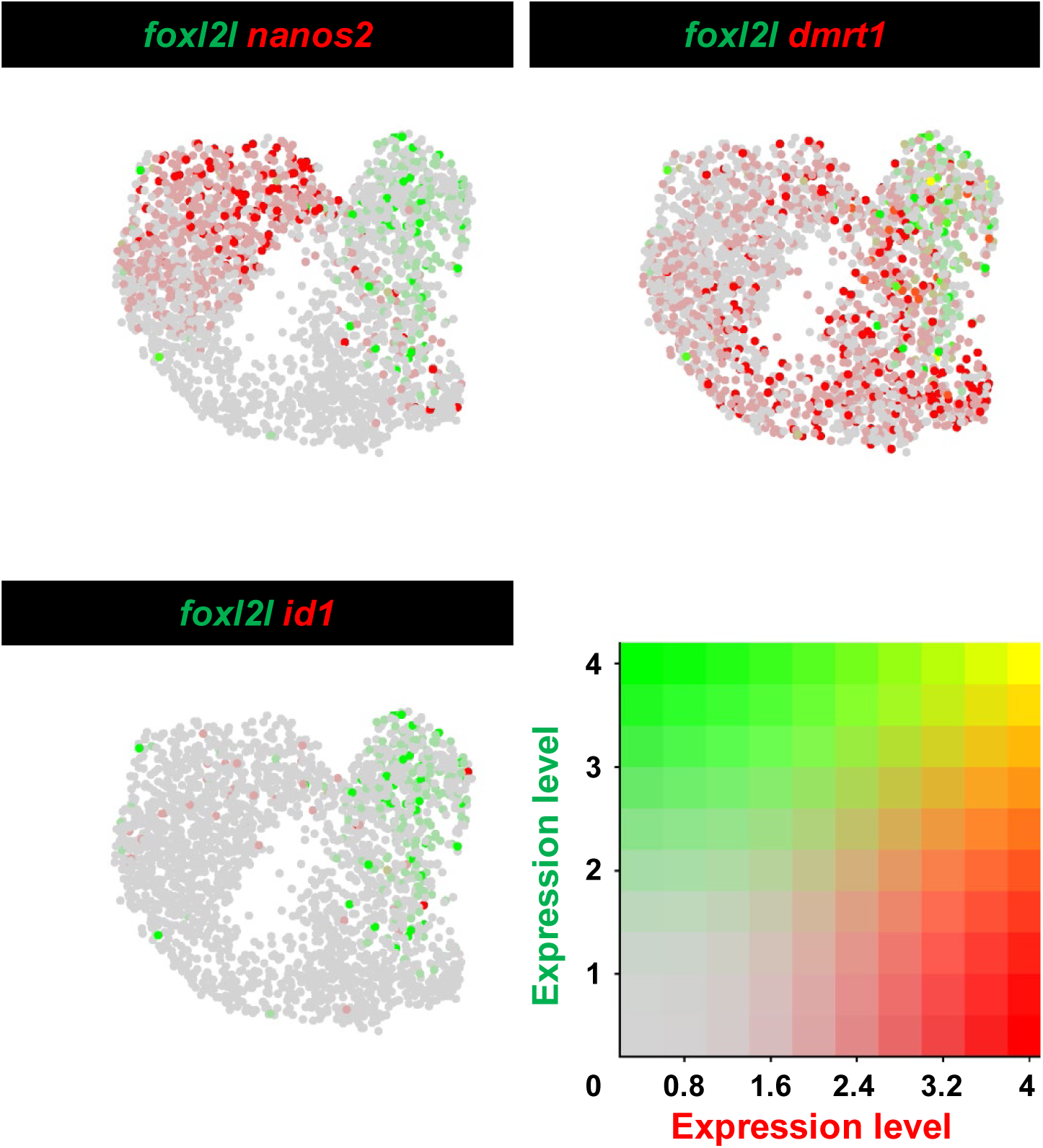
Very few germ cells express *foxl2l* together with *nanos2*, *id1* or *dmrt1* in 40-dpf WT gonad. Expression profile of *foxl2l* with *nanos2*, *id1* or *dmrt1* in 40-dpf germ cell obtained from Y Liu, et al. (Liu et al., 2022) shown in UMAP. The expression values are visualized by colors ranging between 4 to 0.

**Supplementary Fig. S10.**
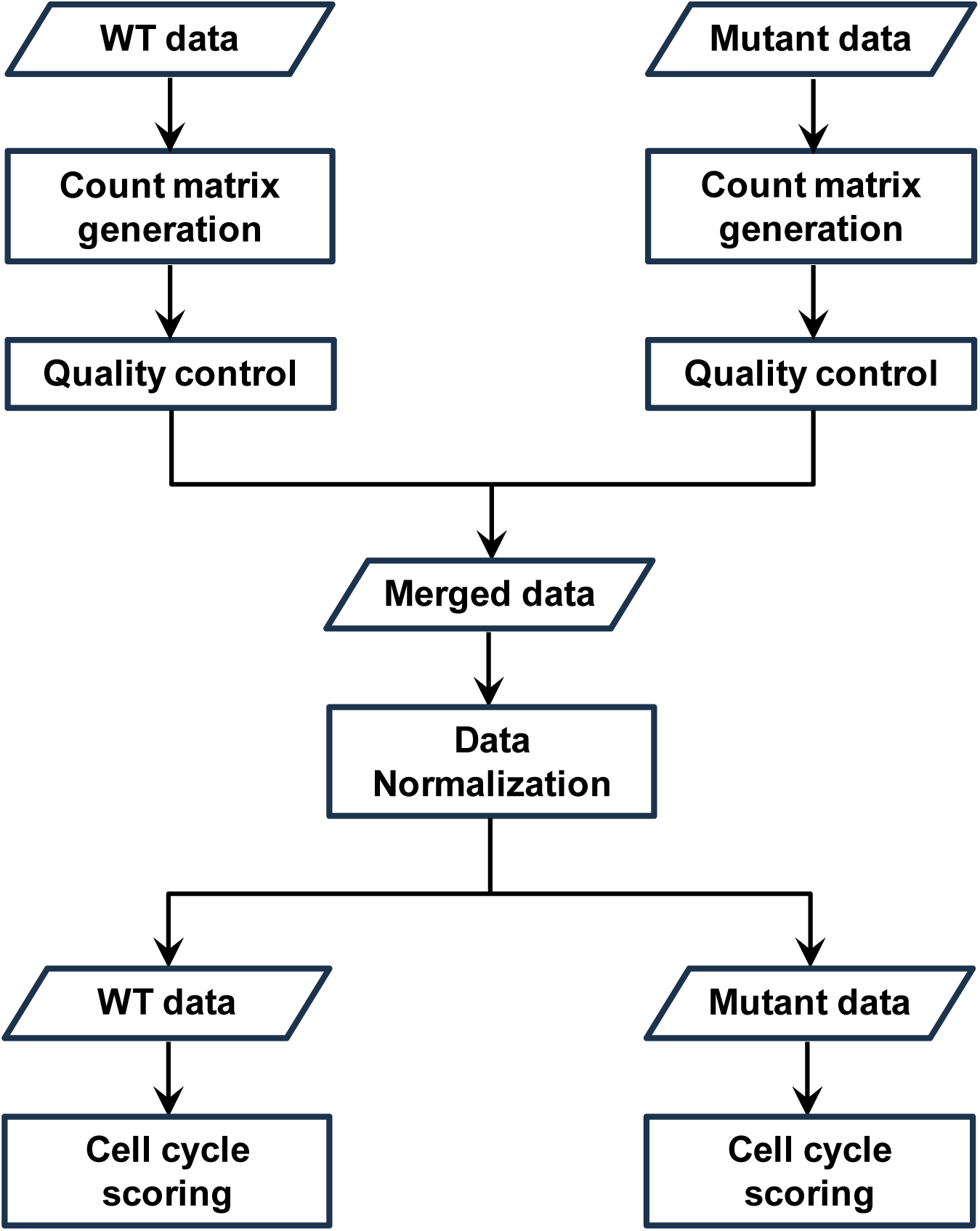
The flow chart of data processing and cell cycle scoring for WT and mutant scRNAseq transcriptome.

**Supplementary Fig. S11.**
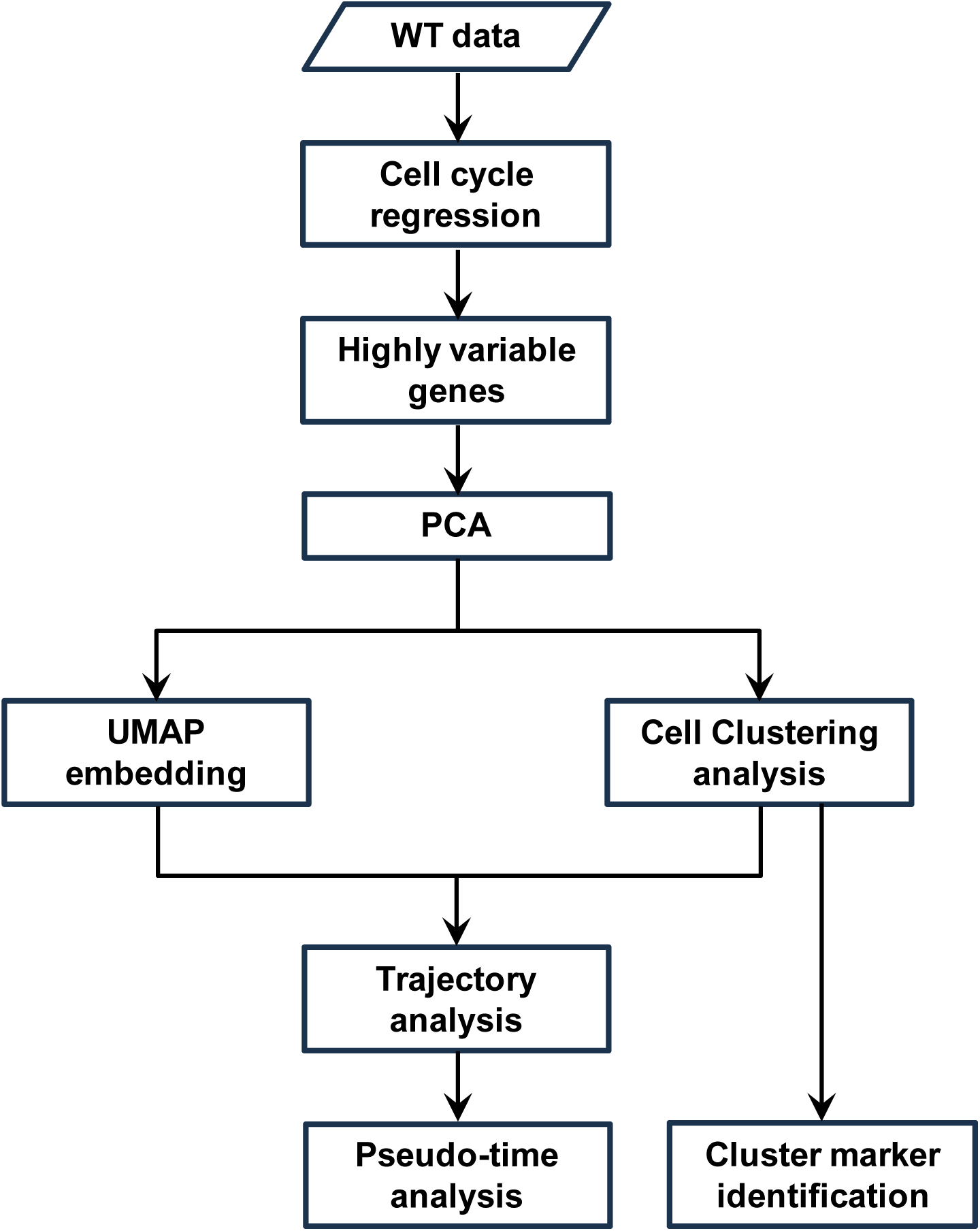
The flow chart of cell clustering analysis for WT cells.

## References

Bais, A. S., & Kostka, D. (2020). scds: computational annotation of doublets in single-cell RNA sequencing data. Bioinformatics, 36(4), 1150–1158. 10.1093/bioinformatics/btz698

Beer, R. L., & Draper, B. W. (2013). nanos3 maintains germline stem cells and expression of the conserved germline stem cell gene nanos2 in the zebrafish ovary. Dev Biol, 374(2), 308–318. 10.1016/j.ydbio.2012.12.003

Benezra, R., Davis, R. L., Lockshon, D., Turner, D. L., & Weintraub, H. (1990). The protein Id: a negative regulator of helix-loop-helix DNA binding proteins. Cell, 61(1), 49–59. 10.1016/0092-8674(90)90214-y

Berta, P., Hawkins, J. R., Sinclair, A. H., Taylor, A., Griffiths, B. L., Goodfellow, P. N., & Fellous, M. (1990a). Genetic-Evidence Equating Sry and the Testis-Determining Factor. Nature, 348(6300), 448–450. Doi 10.1038/348448a0

Berta, P., Hawkins, J. R., Sinclair, A. H., Taylor, A., Griffiths, B. L., Goodfellow, P. N., & Fellous, M. (1990b). Genetic evidence equating SRY and the testis-determining factor. Nature, 348(6300), 448–450. 10.1038/348448A0

Bertho, S., Clapp, M., Banisch, T. U., Bandemer, J., Raz, E., & Marlow, F. L. (2021). Zebrafish dazl regulates cystogenesis and germline stem cell specification during the primordial germ cell to germline stem cell transition. Development, 148(7). 10.1242/dev.187773

Brend, T., & Holley, S. A. (2009). Zebrafish whole mount high-resolution double fluorescent in situ hybridization. J Vis Exp(25). 10.3791/1229

Cao, J., Spielmann, M., Qiu, X., Huang, X., Ibrahim, D. M., Hill, A. J., Zhang, F., Mundlos, S., Christiansen, L., Steemers, F. J., Trapnell, C., & Shendure, J. (2019). The single-cell transcriptional landscape of mammalian organogenesis. Nature, 566(7745), 496–502. 10.1038/s41586-019-0969-x

Cao, Z., Mao, X., & Luo, L. (2019). Germline Stem Cells Drive Ovary Regeneration in Zebrafish. Cell Rep, 26(7), 1709–1717 e1703. 10.1016/j.celrep.2019.01.061

Chen, S. J., Zhang, H. F., Wang, F. H., Zhang, W., & Peng, G. (2016). nr0b1 (DAX1) mutation in zebrafish causes female-to-male sex reversal through abnormal gonadal proliferation and differentiation. Molecular and Cellular Endocrinology, 433(C), 105–116. 10.1016/j.mce.2016.06.005

Dai, S., Qi, S., Wei, X., Liu, X., Li, Y., Zhou, X., Xiao, H., Lu, B., Wang, D., & Li, M. (2021). Germline sexual fate is determined by the antagonistic action of dmrt1 and foxl3/foxl2 in tilapia. Development, 148(8). 10.1242/dev.199380

Dranow, D. B., Hu, K., Bird, A. M., Lawry, S. T., Adams, M. T., Sanchez, A., Amatruda, J. F., & Draper, B. W. (2016). Bmp15 Is an Oocyte-Produced Signal Required for Maintenance of the Adult Female Sexual Phenotype in Zebrafish. PLoS Genet, 12(9), e1006323. 10.1371/journal.pgen.1006323

Gautier, A., Goupil, A. S., Le Gac, F., & Lareyre, J. J. (2013). A promoter fragment of the sycp1 gene is sufficient to drive transgene expression in male and female meiotic germ cells in zebrafish. Biol Reprod, 89(4), 89. 10.1095/biolreprod.113.107706

Ge, C., Ye, J., Zhang, H., Zhang, Y., Sun, W., Sang, Y., Capel, B., & Qian, G. (2017). Dmrt1 induces the male pathway in a turtle species with temperature-dependent sex determination. Development, 144(12), 2222–2233. 10.1242/dev.152033

Hong, S. H., Lee, J. H., Lee, J. B., Ji, J., & Bhatia, M. (2011). ID1 and ID3 represent conserved negative regulators of human embryonic and induced pluripotent stem cell hematopoiesis. J Cell Sci, 124(Pt 9), 1445–1452. 10.1242/jcs.077511

Kantzer, C. G., Yang, W., Grommisch, D., Vikhe Patil, K., Mak, K. H., Shirokova, V., & Genander, M. (2022). ID1 and CEBPA coordinate epidermal progenitor cell differentiation. Development, 149(22). 10.1242/dev.201262

Kikuchi, M., Nishimura, T., Ishishita, S., Matsuda, Y., & Tanaka, M. (2020). foxl3, a sexual switch in germ cells, initiates two independent molecular pathways for commitment to oogenesis in medaka. Proc Natl Acad Sci U S A, 117(22), 12174–12181. 10.1073/pnas.1918556117

Kusz, K. M., Tomczyk, L., Sajek, M., Spik, A., Latos-Bielenska, A., Jedrzejczak, P., Pawelczyk, L., & Jaruzelska, J. (2009). The highly conserved NANOS2 protein: testis-specific expression and significance for the human male reproduction. Mol Hum Reprod, 15(3), 165–171. 10.1093/molehr/gap003

Lin, T., Chao, C., Saito, S., Mazur, S. J., Murphy, M. E., Appella, E., & Xu, Y. (2005). p53 induces differentiation of mouse embryonic stem cells by suppressing Nanog expression. Nat Cell Biol, 7(2), 165–171. 10.1038/ncb1211

Liu, Y., Kossack, M. E., McFaul, M. E., Christensen, L. N., Siebert, S., Wyatt, S. R., Kamei, C. N., Horst, S., Arroyo, N., Drummond, I. A., Juliano, C. E., & Draper, B. W. (2022). Single-cell transcriptome reveals insights into the development and function of the zebrafish ovary. Elife, 11. 10.7554/eLife.76014

Lun, A. T., McCarthy, D. J., & Marioni, J. C. (2016). A step-by-step workflow for low-level analysis of single-cell RNA-seq data with Bioconductor. F1000Res, 5, 2122. 10.12688/f1000research.9501.2

Luzio, A., Santos, D., Monteiro, S. M., & Coimbra, A. M. (2021). Zebrafish male differentiation: Do all testes go through a “juvenile ovary” stage? Tissue Cell, 72, 101545. 10.1016/j.tice.2021.101545

Marlow, F. L., & Mullins, M. C. (2008). Bucky ball functions in Balbiani body assembly and animal-vegetal polarity in the oocyte and follicle cell layer in zebrafish. Dev Biol, 321(1), 40–50. 10.1016/j.ydbio.2008.05.557

Matson, C. K., Murphy, M. W., Sarver, A. L., Griswold, M. D., Bardwell, V. J., & Zarkower, D. (2011). DMRT1 prevents female reprogramming in the postnatal mammalian testis. Nature, 476(7358), 101–104. 10.1038/nature10239

Matsuda, M., Nagahama, Y., Shinomiya, A., Sato, T., Matsuda, C., Kobayashi, T., Morrey, C. E., Shibata, N., Asakawa, S., Shimizu, N., Hori, H., Hamaguchi, S., & Sakaizumi, M. (2002). DMY is a Y-specific DM-domain gene required for male development in the medaka fish. Nature, 417(6888), 559–563. DOI 10.1038/nature751

McConnell, A. M., Yao, C., Yeckes, A. R., Wang, Y., Selvaggio, A. S., Tang, J., Kirsch, D. G., & Stripp, B. R. (2016). p53 Regulates Progenitor Cell Quiescence and Differentiation in the Airway. Cell Rep, 17(9), 2173–2182. 10.1016/j.celrep.2016.11.007

Molchadsky, A., Ezra, O., Amendola, P. G., Krantz, D., Kogan-Sakin, I., Buganim, Y., Rivlin, N., Goldfinger, N., Folgiero, V., Falcioni, R., Sarig, R., & Rotter, V. (2013). p53 is required for brown adipogenic differentiation and has a protective role against diet-induced obesity. Cell Death and Differentiation, 20(5), 774–783. 10.1038/cdd.2013.9

Nüsslein-Volhard, C. a. D., Ralf. (2002). Zebrafish. New York: Oxford University Press.

Nishimura, T., Sato, T., Yamamoto, Y., Watakabe, I., Ohkawa, Y., Suyama, M., Kobayashi, S., & Tanaka, M. (2015). Sex determination. foxl3 is a germ cell-intrinsic factor involved in sperm-egg fate decision in medaka. Science, 349(6245), 328–331. 10.1126/science.aaa2657

Pan, Y. J., Tong, S. K., Hsu, C. W., Weng, J. H., & Chung, B. C. (2022). Zebrafish Establish Female Germ Cell Identity by Advancing Cell Proliferation and Meiosis. Front Cell Dev Biol, 10, 866267. 10.3389/fcell.2022.866267

Qin, M., Zhang, Z., Song, W., Wong, Q. W., Chen, W., Shirgaonkar, N., & Ge, W. (2018). Roles of Figla/figla in Juvenile Ovary Development and Follicle Formation During Zebrafish Gonadogenesis. Endocrinology, 159(11), 3699–3722. 10.1210/en.2018-00648

Sablitzky, F., Moore, A., Bromley, M., Deed, R. W., Newton, J. S., & Norton, J. D. (1998). Stage- and subcellular-specific expression of Id proteins in male germ and Sertoli cells implicates distinctive regulatory roles for Id proteins during meiosis, spermatogenesis, and Sertoli cell function. Cell Growth Differ, 9(12), 1015–1024. https://www.ncbi.nlm.nih.gov/pubmed/9869302

Saito, D., Morinaga, C., Aoki, Y., Nakamura, S., Mitani, H., Furutani-Seiki, M., Kondoh, H., & Tanaka, M. (2007). Proliferation of germ cells during gonadal sex differentiation in medaka: Insights from germ cell-depleted mutant zenzai. Dev Biol, 310(2), 280–290. 10.1016/j.ydbio.2007.07.039

She, Z. Y., & Yang, W. X. (2017). Sry and SoxE genes: How they participate in mammalian sex determination and gonadal development? Semin Cell Dev Biol, 63, 13–22. 10.1016/j.semcdb.2016.07.032

Slanchev, K., Stebler, J., de la Cueva-Mendez, G., & Raz, E. (2005). Development without germ cells: the role of the germ line in zebrafish sex differentiation. Proc Natl Acad Sci U S A, 102(11), 4074–4079. 10.1073/pnas.0407475102

Smith, C. A., Roeszler, K. N., Ohnesorg, T., Cummins, D. M., Farlie, P. G., Doran, T. J., & Sinclair, A. H. (2009). The avian Z-linked gene DMRT1 is required for male sex determination in the chicken. Nature, 461(7261), 267–271. 10.1038/nature08298

Stuart, T., Butler, A., Hoffman, P., Hafemeister, C., Papalexi, E., Mauck, W. M., 3rd, Hao, Y., Stoeckius, M., Smibert, P., & Satija, R. (2019). Comprehensive Integration of Single-Cell Data. Cell, 177(7), 1888–1902 e1821. 10.1016/j.cell.2019.05.031

Suzuki, A., & Saga, Y. (2008). Nanos2 suppresses meiosis and promotes male germ cell differentiation. Genes Dev, 22(4), 430–435. 10.1101/gad.1612708

Thisse, C., & Thisse, B. (2008). High-resolution in situ hybridization to whole-mount zebrafish embryos. Nat Protoc, 3(1), 59–69. 10.1038/nprot.2007.514

Tong, S. K., Hsu, H. J., & Chung, B. C. (2010). Zebrafish monosex population reveals female dominance in sex determination and earliest events of gonad differentiation. Dev Biol, 344(2), 849–856. 10.1016/j.ydbio.2010.05.515

Tsuda, M., Sasaoka, Y., Kiso, M., Abe, K., Haraguchi, S., Kobayashi, S., & Saga, Y. (2003). Conserved role of nanos proteins in germ cell development. Science, 301(5637), 1239–1241. 10.1126/science.1085222

Uchida, D., Yamashita, M., Kitano, T., & Iguchi, T. (2002). Oocyte apoptosis during the transition from ovary-like tissue to testes during sex differentiation of juvenile zebrafish. J Exp Biol, 205(Pt 6), 711–718. 10.1242/jeb.205.6.711

Wang, M., Liu, X., Chang, G., Chen, Y., An, G., Yan, L., Gao, S., Xu, Y., Cui, Y., Dong, J., Chen, Y., Fan, X., Hu, Y., Song, K., Zhu, X., Gao, Y., Yao, Z., Bian, S., Hou, Y., Lu, J., Wang, R., Fan, Y., Lian, Y., Tang, W., Wang, Y., Liu, J., Zhao, L., Wang, L., Liu, Z., Yuan, R., Shi, Y., Hu, B., Ren, X., Tang, F., Zhao, X. Y., & Qiao, J. (2018). Single-Cell RNA Sequencing Analysis Reveals Sequential Cell Fate Transition during Human Spermatogenesis. Cell Stem Cell, 23(4), 599–614 e594. 10.1016/j.stem.2018.08.007

Wassarman, P. M., & Litscher, E. S. (2021). Zona Pellucida Genes and Proteins: Essential Players in Mammalian Oogenesis and Fertility. Genes (Basel), 12(8). 10.3390/genes12081266

Webster, K. A., Schach, U., Ordaz, A., Steinfeld, J. S., Draper, B. W., & Siegfried, K. R. (2017). Dmrt1 is necessary for male sexual development in zebrafish. Dev Biol, 422(1), 33–46. 10.1016/j.ydbio.2016.12.008

Westerfield, M. (2000). The zebrafish book. A guide for the laboratory use of zebrafish (Danio rerio). 4th ed., Univ. of Oregon Press, Eugene.

Wilson, C. A., High, S. K., McCluskey, B. M., Amores, A., Yan, Y. L., Titus, T. A., Anderson, J. L., Batzel, P., Carvan, M. J., 3rd, Schartl, M., & Postlethwait, J. H. (2014). Wild sex in zebrafish: loss of the natural sex determinant in domesticated strains. Genetics, 198(3), 1291–1308. 10.1534/genetics.114.169284

Ying, Q. L., Nichols, J., Chambers, I., & Smith, A. (2003). BMP induction of Id proteins suppresses differentiation and sustains embryonic stem cell self-renewal in collaboration with STAT3. Cell, 115(3), 281–292. 10.1016/s0092-8674(03)00847-x

Yoshida, K., Kondoh, G., Matsuda, Y., Habu, T., Nishimune, Y., & Morita, T. (1998). The mouse RecA-like gene Dmc1 is required for homologous chromosome synapsis during meiosis. Mol Cell, 1(5), 707–718. 10.1016/s1097-2765(00)80070-2

Zhang, N., Yantiss, R. K., Nam, H. S., Chin, Y., Zhou, X. K., Scherl, E. J., Bosworth, B. P., Subbaramaiah, K., Dannenberg, A. J., & Benezra, R. (2014). ID1 is a functional marker for intestinal stem and progenitor cells required for normal response to injury. Stem Cell Reports, 3(5), 716–724. 10.1016/j.stemcr.2014.09.012

Zheng, G. X., Terry, J. M., Belgrader, P., Ryvkin, P., Bent, Z. W., Wilson, R., Ziraldo, S. B., Wheeler, T. D., McDermott, G. P., Zhu, J., Gregory, M. T., Shuga, J., Montesclaros, L., Underwood, J. G., Masquelier, D. A., Nishimura, S. Y., Schnall-Levin, M., Wyatt, P. W., Hindson, C. M., Bharadwaj, R., Wong, A., Ness, K. D., Beppu, L. W., Deeg, H. J., McFarland, C., Loeb, K. R., Valente, W. J., Ericson, N. G., Stevens, E. A., Radich, J. P., Mikkelsen, T. S., Hindson, B. J., & Bielas, J. H. (2017). Massively parallel digital transcriptional profiling of single cells. Nat Commun, 8, 14049. 10.1038/ncomms14049

## References

Liu, Y., Kossack, M. E., McFaul, M. E., Christensen, L. N., Siebert, S., Wyatt, S. R., . . . Draper, B. W. (2022). Single-cell transcriptome reveals insights into the development and function of the zebrafish ovary. Elife, 11. doi:10.7554/eLife.76014

